# A Stu2-mediated intrinsic tension-sensing pathway promotes chromosome biorientation in vivo

**DOI:** 10.1101/453811

**Authors:** Matthew P. Miller, Rena K. Evans, Alex Zelter, Elisabeth A. Geyer, Michael J. MacCoss, Luke M. Rice, Trisha N. Davis, Charles L. Asbury, Sue Biggins

## Abstract

Accurate segregation of chromosomes to daughter cells is a critical aspect of cell division. It requires the kinetochores on duplicated chromosomes to biorient, attaching to microtubules from opposite poles of the cell. Bioriented attachments come under tension, while incorrect attachments lack tension and must be destabilized. A well-studied error correction pathway is mediated by the Aurora B kinase, which destabilizes low tension-bearing attachments. We recently discovered that in vitro, kinetochores display an additional intrinsic tension-sensing pathway that utilizes Stu2. This pathway’s contribution to error correction in cells, however, was unknown. Here, we identify a Stu2 mutant that abolishes its kinetochore function and show that it causes error correction defects in vivo. We also show that this intrinsic tension-sensing pathway functions in concert with the Aurora B-mediated pathway. Together, our work indicates that cells employ at least two pathways to ensure biorientation and the accuracy of chromosome segregation.

## INTRODUCTION

Faithful partitioning of the genetic material is a fundamental aspect of cell division. The maintenance of cellular and organismal fitness requires this process to be executed precisely for every pair of chromosomes. Chromosome segregation errors are the most prevalent genetic alteration in tumor cells and are proposed to be a major factor in the evolution of cancer (reviewed in (Targa and Rancati, 2018). Segregation is mediated by the kinetochore, a highly conserved protein complex that physically attaches chromosomes to the spindle microtubules that ultimately pull the chromosomes apart in anaphase. To ensure that the duplicated chromosomes become segregated to each daughter cell, sister kinetochores must attach to microtubules from opposite poles, a state known as biorientation. Once kinetochores biorient, they come under tension from opposing microtubule-pulling forces. For this process to operate faithfully, kinetochores must “sense” whether proper bioriented microtubule attachments have been made and correct erroneous attachments (reviewed in (Lampson and Grishchuk, 2017)). Pioneering work has yielded insight into the underlying mechanism by showing that incorrect kinetochore attachments are unstable due to the absence of tension (Nicklas and Koch, 1969). Thus, it is the selective release of attachments lacking tension that gives the cell another chance to establish proper attachments.

The canonical “error correction” pathway is mediated by the conserved protein kinase Aurora B. The activity of Aurora B destabilizes kinetochore-microtubule interactions that are not under tension by phosphorylating multiple outer kinetochore components (Biggins et al., 1999; Cheeseman et al., 2002; DeLuca et al., 2006; Hauf et al., 2003; Tanaka et al., 2002). However, it is unclear whether this Aurora B-mediated pathway is solely responsible for correcting erroneous attachments. Recently, we discovered another pathway implicated in the correction of erroneous kinetochore-microtubule attachments (Akiyoshi et al., 2010; Miller et al., 2016). Using a reconstitution system, we found (i) that kinetochores exhibit an intrinsic selectivity for high tension-bearing attachments that is independent of Aurora B (Akiyoshi et al., 2010), and (ii) that this direct mechano-sensitivity depends on the kinetochore-associated activity of the XMAP215 family member, Stu2 (Miller et al., 2016). Kinetochore-associated Stu2 enables long-lived attachments, especially when tension is high, by preventing detachment specifically during microtubule assembly. Conversely, Stu2 helps ensure that kinetochore attachments lacking tension are short-lived by promoting detachment during microtubule disassembly at low force. These combined effects result in selective stabilization of high tension-bearing kinetochore attachments in vitro (Miller et al., 2016). However, the extent to which this intrinsic tension-sensing pathway regulates kinetochore-microtubule attachments in vivo, as well as how it integrates with the Aurora B pathway, has yet to be determined.

Here, we address the function of Stu2 at kinetochores in vivo to analyze the role of the intrinsic tension-sensing pathway in cells. Stu2 performs numerous cellular functions (reviewed in (Al-Bassam and Chang, 2011), so we looked for Stu2 mutations that abolish its kinetochore localization but that maintain normal spindle length. We found that a Stu2 mutant lacking the coiled-coil domain cannot associate with kinetochores or enhance kinetochore attachment stability in vitro. When expressed in cells, this mutant results in defective kinetochore biorientation and leads to a spindle checkpoint-dependent cell cycle delay. Furthermore, Stu2’s kinetochore function is necessary for the establishment of bioriented attachments but is dispensable thereafter, reminiscent of the requirement of Aurora B in error correction. Finally, kinetochore-associated Stu2 acts in concert with Aurora B; perturbing both pathways results in an additive growth defect and an increased rate of chromosome missegregation. Together, our findings suggest that mitotic error correction in vivo requires both the Aurora B-mediated and Stu2-dependent intrinsic tension-sensing pathways. Our results provide further mechanistic insight into the process of error correction as well as the manner in which tension promotes accurate chromosome segregation.

## RESULTS

### Dimerization-deficient mutant of Stu2 is defective in kinetochore association yet supports normal mitotic spindle formation

Stu2 has important roles in both the formation of the mitotic spindle and the regulation of kinetochore function (Al-Bassam et al., 2006; Kosco et al., 2001; Miller et al., 2016; Pearson et al., 2003). With the goal of identifying a mutant that disrupts Stu2’s kinetochore localization but that supports normal mitotic spindle formation, we set out to 1) map the domain(s) required for its kinetochore association, and 2) examine the spindle length in cells expressing these mutants. XMAP215 family members are large proteins that contain variable numbers of highly conserved tumor over-expressed gene (TOG) domain arrays that bind to curved tubulin dimers (Al-Bassam et al., 2006; Ayaz et al., 2012, 2014; Fox et al., 2014) as well as a basic linker domain that promotes binding to the microtubule lattice (Geyer et al., 2018; Wang and Huffaker, 1997). The budding yeast Stu2 protein contains two N-terminal TOG domains, followed by a basic linker domain (SK-rich), a homo-dimerization domain (coiled-coil), and a small C-terminal domain that is required for Stu2’s association with the microtubule-associated proteins (MAPs) Bik1, Bim1 and Spc72 (Figure 1A; (Al-Bassam et al., 2006; Usui et al., 2003; Wang and Huffaker, 1997; Wolyniak et al., 2006)).

**Figure 1.**
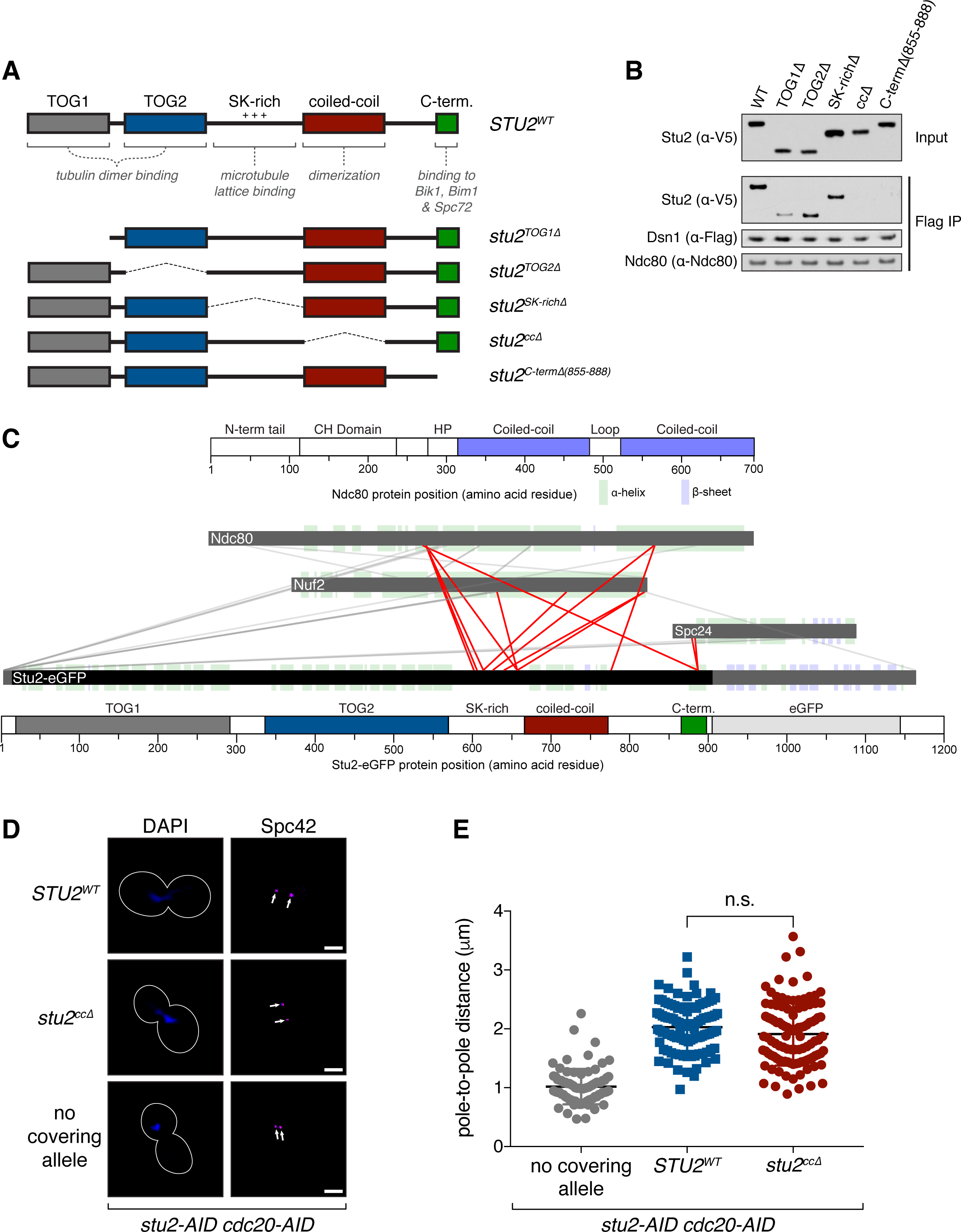
Dimerization-deficient mutant of Stu2 is defective in kinetochore association yet supports normal mitotic spindle formation. A) Schematic of Stu2’s domain architecture and corresponding deletions. Note: Internal deletions (i.e. not at the N- or C-terminus) were made by inserting a linker, denoted by a dashed line. B) Exponentially growing *stu2-AID* cultures expressing an ectopic copy of *STU2* (*STU2^WT^*, SBY13901; *stu2^TOG1Δ^*, SBY13904; *stu2^TOG2Δ^*, SBY13907; *stu2^SK-richΔ^*, SBY13913; *stu2^ccΔ^*, SBY13916; or *stu2^C-termΔ(855-888)^*, SBY14269) that also contained Dsn1-6His-3Flag were treated with auxin 30 min prior to harvesting. Protein lysates were subsequently prepared and kinetochore particles were purified by α-Flag immunoprecipitation (IP) and analyzed by immunoblotting. C) Cross-linking mass spectrometry analysis reveals interactions between recombinant Stu2-eGFP and the Ndc80 complex. Cross-links formed with EDC between Stu2 and the Ndc80 complex are depicted by red lines. Bar diagrams with structural features of Ndc80 and Stu2-eGFP proteins are included. (CH: calponin homology; HP: hairpin; TOG: tumor overexpressed gene; SK-rich: regions with stretches of sequences rich in Serine, Glycine, Lysine). For clarity, cross-links within Ndc80 complex proteins and those involving Stu2 fusion regions (extreme N-terminus and eGFP) are shown in grey. Data are shown for peptides with Percolator assigned q-values ≤ 0.05 and a minimum of 2 PSMs. D) Exponentially growing *stu2-AID cdc20-AID* cultures with an ectopically expressed *STU2* allele (*STU2^WT^*, SBY17369; *stu2^ccΔ^*, SBY17371) or without an ectopic allele (no covering allele, SBY17367) that also contained *MTW1-3GFP* (kinetochore) and *SPC42-CFP* (spindle pole; marked with white arrows) were treated with auxin for 2 h to arrest cells in metaphase. Representative images for each are shown. White bars represent 2 µm. E) Spindle length (spindle pole-to-pole distance) and kinetochore distribution (distance between bi-lobed kinetochore clusters; Figure 4C) was measured for cells described in (D). n = 80-105 cells; p values were determined using a two-tailed unpaired t test (n.s. = not significant).

To determine the element(s) within Stu2 required for kinetochore-association, we individually deleted each of these domains and expressed the mutant alleles from an ectopic locus in a strain harboring a conditional *stu2-AID* allele at the endogenous locus (Miller et al., 2016). Each of these mutant alleles was expressed at or near wild-type levels (Figure 1B & Figure S1E, inputs). The localization of a similar set of *stu2* mutations was recently examined in cells (Humphrey et al., 2018), but only in the presence of wild-type endogenous protein, which through homo-dimerization and/or competition could affect the localization of the mutant proteins. We therefore sought to examine the mutants in the absence of endogenous protein. While we attempted to examine their kinetochore localization in cells using GFP-tagged alleles, distinguishing the pool of Stu2 on the microtubule tip from that on the kinetochore made confidently assaying the sub-cellular localization of the mutant proteins too difficult (data not shown). We therefore analyzed the levels of the Stu2 mutants on purified kinetochores by immunoblotting. We depleted the endogenous Stu2-AID protein by adding auxin and then isolated native kinetochores via single-step immunoprecipitation of the Mis12/MIND/Mtw1 complex component Dsn1-His-Flag (Akiyoshi et al., 2010). While most of the mutations did not affect the Stu2-kinetochore interaction, deletion of either the coiled-coil or C-terminal domains abolished Stu2’s ability to co-purify with native kinetochores (Figure 1B & Figure S1E). Deletion of the TOG1 domain reduced but did not abolish Stu2’s kinetochore interaction. To further confirm the requirement of these domains we used in vitro binding assays. We purified and immobilized kinetochores lacking endogenous Stu2 and examined the ability of each Stu2 variant (purified independently) to bind the immobilized kinetochores. Consistent with our co-purification results, deletions of the coiled-coil, C-terminal, and TOG1 domains disrupted the ability of purified Stu2 to ‘re-bind’ the kinetochores (Figure S1A-D). These in vitro binding data are consistent with the localization data of similar mutants analyzed in cells (Humphrey et al., 2018).

As a parallel approach to define regions of Stu2 required for its kinetochore interaction, we conducted crosslinking mass spectrometry with recombinant Stu2-eGFP and the outer kinetochore Ndc80 complex (Ndc80c), which is Stu2’s kinetochore receptor (Figure S1F; (Hsu and Toda, 2011; Miller et al., 2016; Tang et al., 2013). We observed crosslinks within the heterotetrameric Ndc80c (consistent with previous reports; (Kim et al., 2017; Tien et al., 2014)), and also numerous crosslinks between Ndc80c and Stu2. We identified several crosslinks between the C-terminal domain of Stu2 and Ndc80c, consistent with our observation that the C-terminus is required for kinetochore association (Figure 1C). Interestingly, even though Stu2’s coiled-coil domain is required for its kinetochore association, we did not detect any crosslinks between this domain and Ndc80c. This observation suggests that Stu2’s coiled-coil domain may not directly mediate binding to the Ndc80c; instead, Stu2’s kinetochore-association may require homo-dimerization of the protein. Unexpectedly, we also detected numerous crosslinks between Stu2’s basic linker (SK-rich) domain and Ndc80c even though this domain is dispensable for Stu2-Ndc80c binding (Figure 1B-C & S1F). In summary, we find that the coiled-coil and C-terminal domains of Stu2 are required for its association with kinetochores (a summary of in vitro binding data can be found in Figure S1G).

Stu2 is a microtubule polymerase that has important roles in the formation of the mitotic spindle, which has made it difficult to analyze its kinetochore function in cells (Al-Bassam et al., 2006; Kosco et al., 2001; Pearson et al., 2003). We therefore sought after a Stu2 mutant that lacked kinetochore association but maintained spindle functions. While both the coiled-coil and C-terminal domains are required for Stu2’s kinetochore association, the C-terminal domain has established interactions with proteins involved in microtubule function (Stangier et al., 2018; Usui et al., 2003; Wolyniak et al., 2006). We therefore tested whether the *stu2* mutant lacking the coiled-coil domain (henceforth referred to as *stu2^cc∆^*) had normal spindle length. We arrested cells in metaphase by depleting Cdc20 (using a *cdc20-AID* allele) and examined the distance between spindle poles (labeled by Spc42-CFP). The cells also carried a *stu2-AID* allele to deplete the endogenous Stu2-AID protein and an ectopic copy of the *STU2* variant to be examined. As expected, cells that did not express a covering *STU2* allele (and thus had no Stu2 protein present after degradation of endogenous Stu2-AID), arrested with a short bipolar spindle (1.02 ± 0.30 μm distance between Spc42 foci; (Kosco et al., 2001; Miller et al., 2016; Pearson et al., 2003)), whereas cells that expressed a wild-type copy of *STU2* arrested with a longer and characteristic metaphase-length spindle (2.03 ± 0.40 μm; Figure 1D & 1E). Cells that expressed the *stu2^ccΔ^* allele arrested with a spindle length (1.91 ± 0.54 μm) that was nearly indistinguishable from that of cells expressing a wild-type copy of *STU2*. Thus, while mutants that alter Stu2’s dimerization exhibit polymerization defects in vitro (Al-Bassam et al., 2006; Geyer et al., 2018), our data indicate that this does not lead to a defect in spindle formation in vivo. Furthermore, the fact that the *stu2^ccΔ^* mutant is defective in kinetochore binding yet supports a normal mitotic spindle provides a tool to examine Stu2’s kinetochore functions in cells.

### Homo-dimerization is required for Stu2’s kinetochore association

Although Stu2’s coiled-coil domain is required for its kinetochore association, we did not detect any crosslinks between this domain and Ndc80c (Figure 1C). This suggests that Stu2’s kinetochore-association may simply require the protein to form a homo-dimer, and not sequence specific elements within the coiled-coil domain. To test this, we replaced the coiled-coil domain with a different homo-dimerization domain (that of the bZIP transcriptional activator, Gcn4, to generate *stu2^ccα::bZIP^* (Ellenberger et al., 1992). Restoring the ability of Stu2 to homo-dimerize rescued its kinetochore association (Figure 2A; (Humphrey et al., 2018)). We next examined how disrupting the Stu2-kinetochore interaction affected cell growth by examining the ability of the *stu2^ccΔ^* allele to rescue the lethality resulting from depletion of the endogenous Stu2 (Al-Bassam et al., 2006; Ayaz et al., 2014; Miller et al., 2016; Pearson et al., 2003). We found that the coiled-coil deletion resulted in impaired growth and benomyl sensitivity (Figure 2B & S2B). Similar to our kinetochore localization findings, restoring the ability of Stu2 to homo-dimerize (using the *stu2^ccΔ::bZIP^* allele) fully rescued the growth defects of the coiled-coil deletion. These data suggest that disrupting Stu2’s kinetochore localization leads to a significant growth defect.

**Figure 2.**
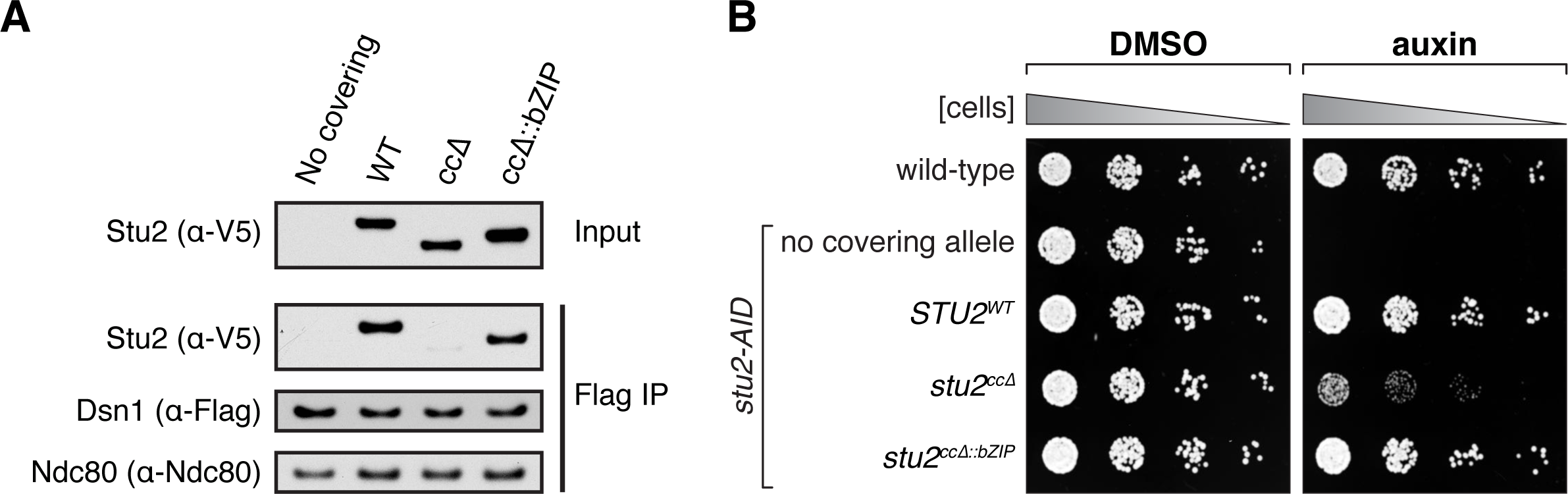
Restoring homo-dimerization rescues Stu2’s kinetochore association. A) Exponentially growing Stu2-AID cultures with an ectopic copy of *STU2* (*STU2^WT^*, SBY13901; *stu2^ccΔ^*, SBY13916; or *stu2^ccΔ::bZIP^*, SBY13935) or without an ectopic allele (no covering allele, SBY13772) that also contained Dsn1-6His-3Flag were treated with auxin 30 min prior to harvesting. Protein lysates were subsequently prepared and kinetochore particles were purified by α-Flag immunoprecipitation (IP) and analyzed by immunoblotting. B) Wild-type (SBY3), *stu2-AID* (no covering allele, SBY13772) and *stu2-AID* cells expressing various *STU2-3V5* alleles from an ectopic locus (*STU2^WT^*, SBY13901; *stu2^ccΔ^*, SBY13916; *stu2^ccΔ::bZIP^*, SBY13935) were serially diluted (5-fold) and spotted on plates containing either DMSO or 500 µM auxin. Refer to Figure S3 for a similar analysis of all alleles examined.

### Kinetochore association is required for Stu2’s ability to enhance attachment stability in vitro

We next tested whether Stu2’s kinetochore association is required for its ability to strengthen Ndc80-based attachments in vitro. We used an optical trapping-based ‘‘force-ramp’’ technique in which native Ndc80c was linked to beads and then attached to growing microtubule tips using a laser trap (Figure 3A; reviewed in (Franck et al., 2010)). We gradually increased the force across the Ndc80c-microtubule interface until the attachment ruptured. When beads contained Ndc80c alone, they formed attachments with an average rupture strength of 3.7 ± 0.3 pN (Miller et al., 2016; Powers et al., 2009). Addition of purified wild-type Stu2 increased the rupture strength of these Ndc80c-based attachments dramatically, to an average of 10.6 ± 0.6 pN (Miller et al., 2016). In contrast, the strength of Ndc80-based attachments was much lower upon addition of the purified, kinetochore-binding deficient Stu2^ccΔ^ mutant, with an average rupture strength of only 5.6 ± 0.6 pN (Figure 3B & 3C). This observation shows that the Stu2^ccΔ^ mutant disrupts a major kinetochore function in vitro, consistent with its defect in kinetochore localization.

**Figure 3.**
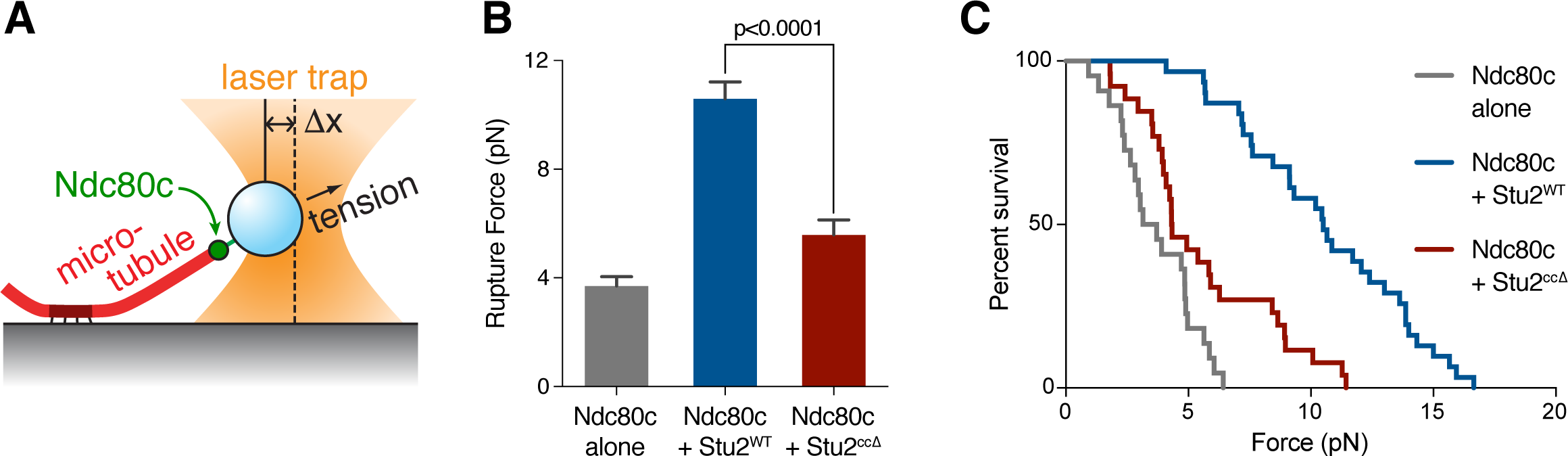
Kinetochore-binding by Stu2 is required for enhancing Ndc80c-microtubule attachment stability. A) Schematic of optical trap assay. Dynamic microtubules are grown from coverslip-anchored seeds. Beads linked to Ndc80c via Spc24 are manipulated using an optical trap to exert applied force across the Ndc80c-microtubule interface in the presence or absence of purified Stu2. B) Mean rupture forces for Ndc80c-linked beads untreated or incubated with Stu2^WT^ or Stu2^ccΔ^. Error bars represent SEM (n = 22–31 events). P value was determined using a two-tailed unpaired t test. All measurements were conducted within the same experimental set, but the data for Ndc80c alone and in the presence of Stu2^WT^ are reported previously in (Miller et al., 2016). C) Attachment survival probability versus force for the data in (B).

### Abolishing Stu2 kinetochore localization results in a kinetochore attachment defect and spindle checkpoint-dependent cell cycle delay

Our identification of a *stu2* mutant that is defective in kinetochore function but that can still promote the formation of a normal mitotic spindle gave us the ability to examine the effects of removing Stu2 from kinetochores in cells. To do this, we first analyzed metaphase kinetochore distribution in cells expressing the dimerization-defective *stu2* mutant. The cells carried a *stu2-AID* allele to allow us to deplete the endogenous Stu2-AID protein and reveal the phenotype of the mutant. Cells were arrested in metaphase by degrading Cdc20 (using a *cdc20-AID* allele) and kinetochore distribution was assayed using Mtw1-3GFP. In budding yeast, properly attached bioriented kinetochores cluster and exhibit a characteristic bi-lobed distribution at metaphase when they come under tension (Goshima and Yanagida, 2000; He et al., 2001; Pearson et al., 2001). As expected, the vast majority of cells (89% ± 1%) expressing a wild-type allele of *STU2* exhibited normal spindle length and a normal bi-lobed kinetochore distribution (Figure 4A & 4B). Consistent with previous observations (Kosco et al., 2001; Marco et al., 2013; Miller et al., 2016; Pearson et al., 2003; Severin et al., 2001), cells depleted of Stu2 had extremely short spindles and three classes of kinetochore configurations: bi-lobed (32% ± 4%), mono-lobed (47% ± 8%), and unattached/off spindle axis (21% ± 4%; (Pinsky et al., 2006). In contrast, the cells expressing the *stu2^ccΔ^* allele had normal length spindles but a significantly altered kinetochore alignment (Figure 1D-E & Figure 4A-B). Although the majority of *stu2^ccΔ^* cells displayed a bi-lobed kinetochore configuration (82% ± 5%), the clusters appeared closer together than *STU2^WT^* and there was a significant number of cells that also displayed unattached kinetochores (12% ± 1%; Figure 4B). We measured the distance between the Mtw1-3GFP foci in cells displaying a bi-lobed kinetochore configuration. The kinetochore foci were 1.23 ± 0.31 μm apart in cells expressing *STU2^WT^* (Figure 4A & 4C). The cells expressing *stu2^ccΔ^* exhibited a significantly shorter distance of 0.89 ± 0.29 μm between kinetochore foci (compared to the 0.38 ± 0.15 μm for cells not expressing a copy of *STU2*; Figure 4A & 4C), suggesting that the kinetochores do not come under normal levels of tension. Together, these results show that abolishing Stu2’s kinetochore localization in vivo supports the formation of a normal length mitotic spindle but leads to defects kinetochore tension and attachments.

**Figure 4.**
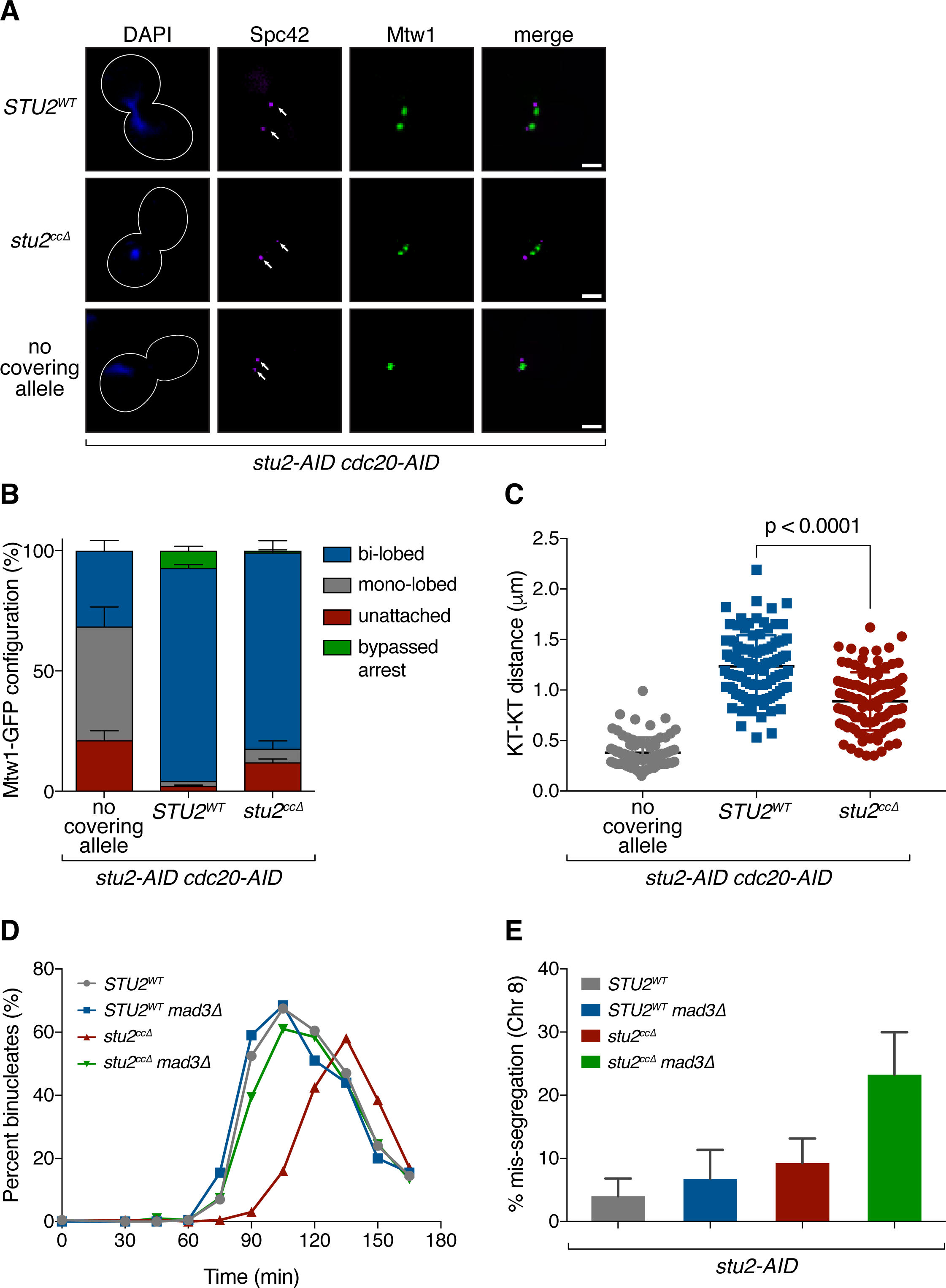
Kinetochore-binding deficient Stu2 exhibits a biorientation defect and spindle checkpoint-dependent cell cycle delay. A) Exponentially growing *stu2-AID cdc20-AID* cultures with an ectopically expressed *STU2* allele (*STU2^WT^*, SBY17369; *stu2^ccΔ^*, SBY17371) or without an ectopic allele (no covering allele, SBY17367) that also contained *MTW1-3GFP* (kinetochore) and *SPC42-CFP* (spindle pole; marked with white arrows) were treated with auxin for 2 h to arrest cells in metaphase. Representative images for each are shown. White bars represent 2 µm. B) Quantification of Mtw1 localization from (A). Error bars represent SD of two independent experiments; n = 100 cells for each experiment. C) Kinetochore distribution (distance between bi-lobed kinetochore clusters) and spindle length (spindle pole-to-pole distance; Figure 1E) was measured for cells described in (A). n = 80-105 cells; p values were determined using a two-tailed unpaired t test. D) *stu2-AID* cells with an ectopically expressed *STU2* allele with and without a spindle checkpoint mutation (*STU2^WT^ MAD3*, SBY17527; *stu2^ccΔ^ MAD3*, SBY17560; *STU2^WT^ mad3Δ*, SBY17668; *stu2^ccΔ^ mad3Δ*, SBY17669] that also contained a fluorescently labeled *CEN8* were released from a G1 arrest into auxin containing media. Cell cycle progression determined by the accumulation of binucleate cells. Shown is a representative experiment. E) Quantification of chromosome mis-segregation in anaphase (percent of binucleate cells with a fluorescently labeled *CEN8* signal in only one of the two nuclei) from (D). Error bars represent SD of two independent experiments; n = 200 cells for each experiment.

Defects in kinetochore-microtubule interactions trigger the spindle checkpoint and result in anaphase onset delays (reviewed in (Lischetti and Nilsson, 2015). To determine if kinetochore alignment defects in the *stu2^ccΔ^* expressing cells delay the cell cycle, we examined cell cycle progression and chromosome segregation. Cells were arrested in G1 and released into auxin-containing media to degrade the endogenous Stu2-AID protein. As above, these cells ectopically expressed either *STU2^WT^* or *stu2^ccΔ^*, and also contained a fluorescently marked centromere on chromosome VIII. Cells expressing *stu2^ccΔ^* displayed an anaphase onset delay (Figure 4D & S4A-E), suggesting spindle checkpoint activation. Consistent with this delay being caused by kinetochore attachment defects, deletion of the spindle checkpoint component *MAD3* suppressed the cell cycle delay and allowed the cells to progress into anaphase (Figure 4D). We analyzed the frequency of mis-segregation of chromosome VIII in these cells and found it was notably increased in *stu2^ccΔ^ mad3Δ* cells (*STU2^WT^* MAD3, 4.0 ± 2.8%; *STU2^WT^* mad3Δ, 6.8 ± 4.6%; *stu2^ccΔ^* MAD3, 9.3 ± 3.9%; *stu2^ccΔ^* mad3Δ, 23.3 ± 4.8%; Figure 4E). Taken together, these data suggest that Stu2 localization to kinetochores promotes correct kinetochore attachments in vivo.

### Kinetochore-associated Stu2 is required for the establishment of bioriented attachments but dispensable for their maintenance

Tension sensing is critical for the establishment of correctly bioriented attachments (reviewed in (Lampson and Grishchuk, 2017). We previously found that Stu2 is required for the tension-dependent stabilization of microtubule attachments in vitro (Miller et al., 2016), potentially implicating this intrinsic tension sensing pathway in the process of biorientation. Our observation that cells expressing the kinetochore-binding deficient *stu2^ccΔ^* allele display abnormal kinetochore clustering (Figure 4A-C) is consistent with defects in the ability to correctly biorient and come under the proper level of tension. To directly examine whether there was a biorientation defect in these cells, we fluorescently marked the centromere of chromosome VIII (Straight et al., 1996), and followed a similar strategy as above: arresting cells in metaphase by depleting Cdc20 and the endogenous Stu2-AID protein, and ectopically expressing either *STU2^WT^*, *stu2^ccΔ^*, or no covering allele. In metaphase, sister kinetochores become bioriented and are transiently stretched apart by opposing microtubule pulling forces (Pearson et al., 2001). In metaphase cells expressing a wild-type copy of *STU2*, 63.0 ± 6% displayed bioriented attachments, as judged by the fluorescently marked centromeres appearing as two distinct foci (Figure 5A). As expected, cells that did not express a covering allele of *STU2* had only 6.0 ± 1% of cells with bioriented attachments. We found that cells expressing the *stu2^ccΔ^* allele also displayed a substantial biorientation defect, with only 15.5 ± 2% of cells displaying two distinct centromere-marked foci. These findings indicate that the Stu2-dependent intrinsic tension sensing pathway plays an important role in the *establishment* of bioriented attachments in vivo, similar to the canonical Aurora B-mediated error correction pathway.

**Figure 5.**
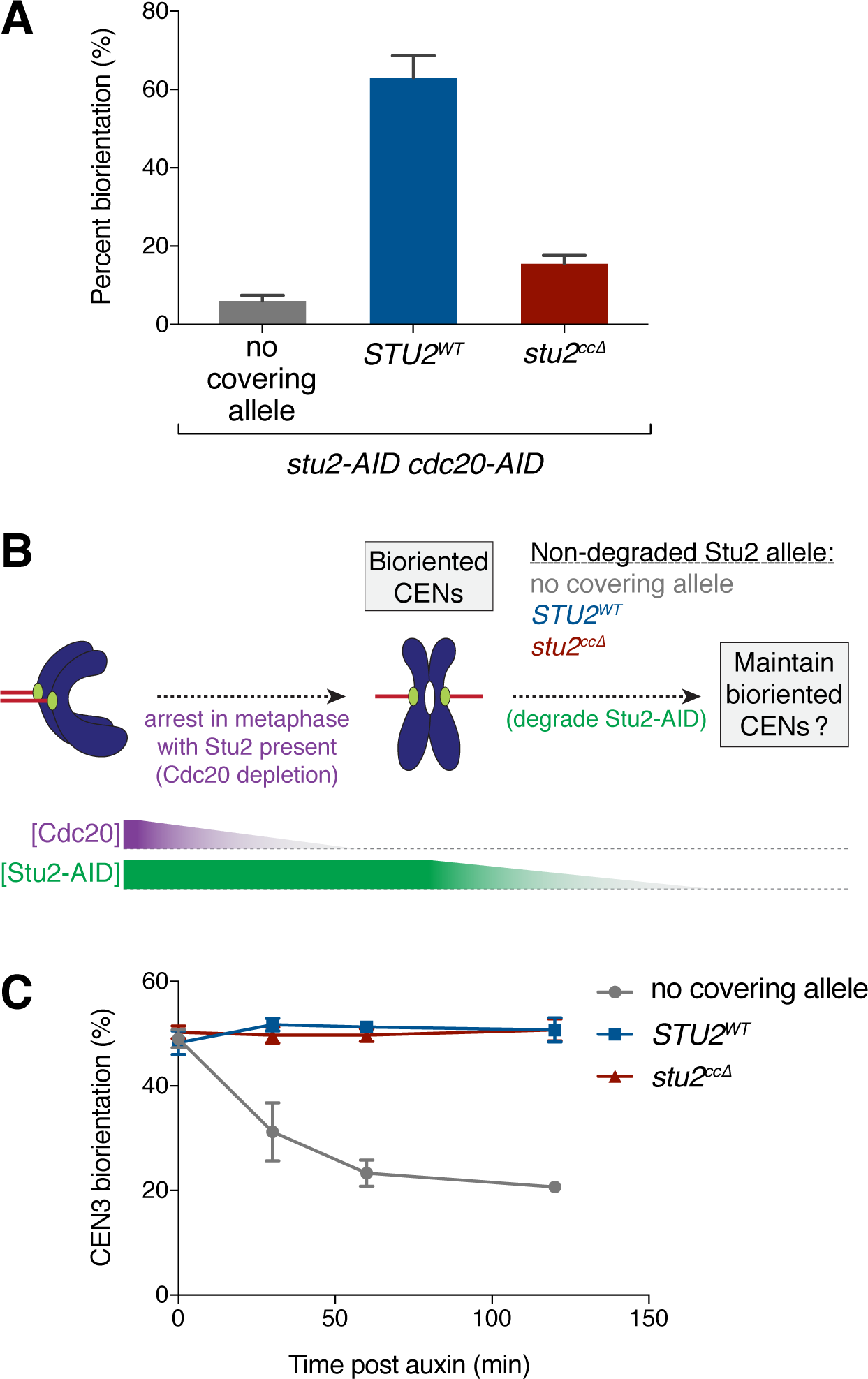
Kinetochore-associated Stu2 is not required for maintenance of biorientation. A) Exponentially growing *stu2-AID cdc20-AID* cultures with an ectopically expressed *STU2* allele (*STU2^WT^*, SBY17748; *stu2^ccΔ^*, SBY17750) or without an ectopic allele (no covering allele, SBY17708) that also contained a fluorescently labeled *CEN8* were treated with auxin for 2 h to arrest cells in metaphase. Percentage of cells that display two distinct GFP foci (i.e. bioriented *CEN8*) was quantified. Shown is the average of two independent experiments, error bars represent SD (n ≧ 100 cells). B) Schematic of experiment in (C). C) Exponentially growing *stu2-AID pMET-CDC20* cells that contained a fluorescently labeled *CEN3* and an ectopically expressed *STU2* allele (*STU2^WT^*, SBY18370; *stu2^ccΔ^*, SBY18371) or without an ectopic allele (no covering allele, SBY18359) were arrested in metaphase by the addition of methionine for 3 h. Once arrested in metaphase, auxin was added to degrade the Stu2-AID protein and the percentage of cells that display two distinct GFP foci (i.e. bioriented *CEN3*) was quantified over time. Error bars represent SD of three independent experiments; n = 100 cells for each time point.

Aurora B function is dispensable once bioriented attachments have been made (Tanaka et al., 2002), so we tested whether this was also the case for the Stu2-mediated pathway. To assay the maintenance of biorientation, we arrested *stu2-AID* cells containing a fluorescently marked *CENIII* in metaphase to establish biorientation. For these experiments, we used a methionine repressible *pMET-CDC20* allele to achieve the metaphase arrest. All cells displayed a similar level of biorientation (Figure 5C). We then added auxin to deplete the endogenous Stu2-AID protein and analyzed the maintenance of *CEN3* biorientation. As expected, cells expressing *STU2^WT^* maintained bioriented attachments after the addition of auxin (50.7 ± 2.3% at 2 h post auxin addition; Figure 5C). In cells without a covering *STU2* allele, however, the number of cells with bioriented attachments quickly decreased after the addition of auxin (20.7 ± 0.6% at 2 h post auxin), likely due to mitotic spindle collapse (Figure S5). In contrast, in cells expressing the *stu2^ccΔ^* allele, the level of biorientation was maintained after the addition of auxin and appeared indistinguishable from that of wild-type (50.7 ± 2.1%). These results suggest that Stu2’s kinetochore-associated function, and by inference the intrinsic mechanical tension-sensing pathway that it mediates, is only required for the *establishment* of correctly bioriented attachments and is dispensable thereafter.

### Kinetochore localization-deficient mutant of Stu2 displays synthetic phenotype with an Aurora B mutant

Our work suggests that Stu2 promotes kinetochore biorientation, so we tested its contribution relative to the Aurora B-mediated error correction pathway. First, we examined whether perturbing Stu2 kinetochore localization enhanced the defects of an Aurora B mutant using the *stu2^ccΔ^* mutant and the *ipl1-321* temperature sensitive allele of Aurora B (Biggins and Murray, 2001; Biggins et al., 1999). Since Aurora B activity is essential for cell viability, we used a semi-permissive temperature in which *ipl1-321* cells grow nearly as well as wild-type cells. We found that under conditions where both pathways were perturbed (i.e. in the presence of auxin and at the semi-permissive temperature of 30°C), the cells expressing the *stu2^ccΔ^* allele and harboring *ipl1-321* displayed a significant additive growth defect over either of the single mutations (Figure 6A & S6). We also observed a similar additive growth defect for the double mutant when grown at the permissive temperature (23°C) but in the presence of low concentrations of benomyl (Figure 6A).

**Figure 6.**
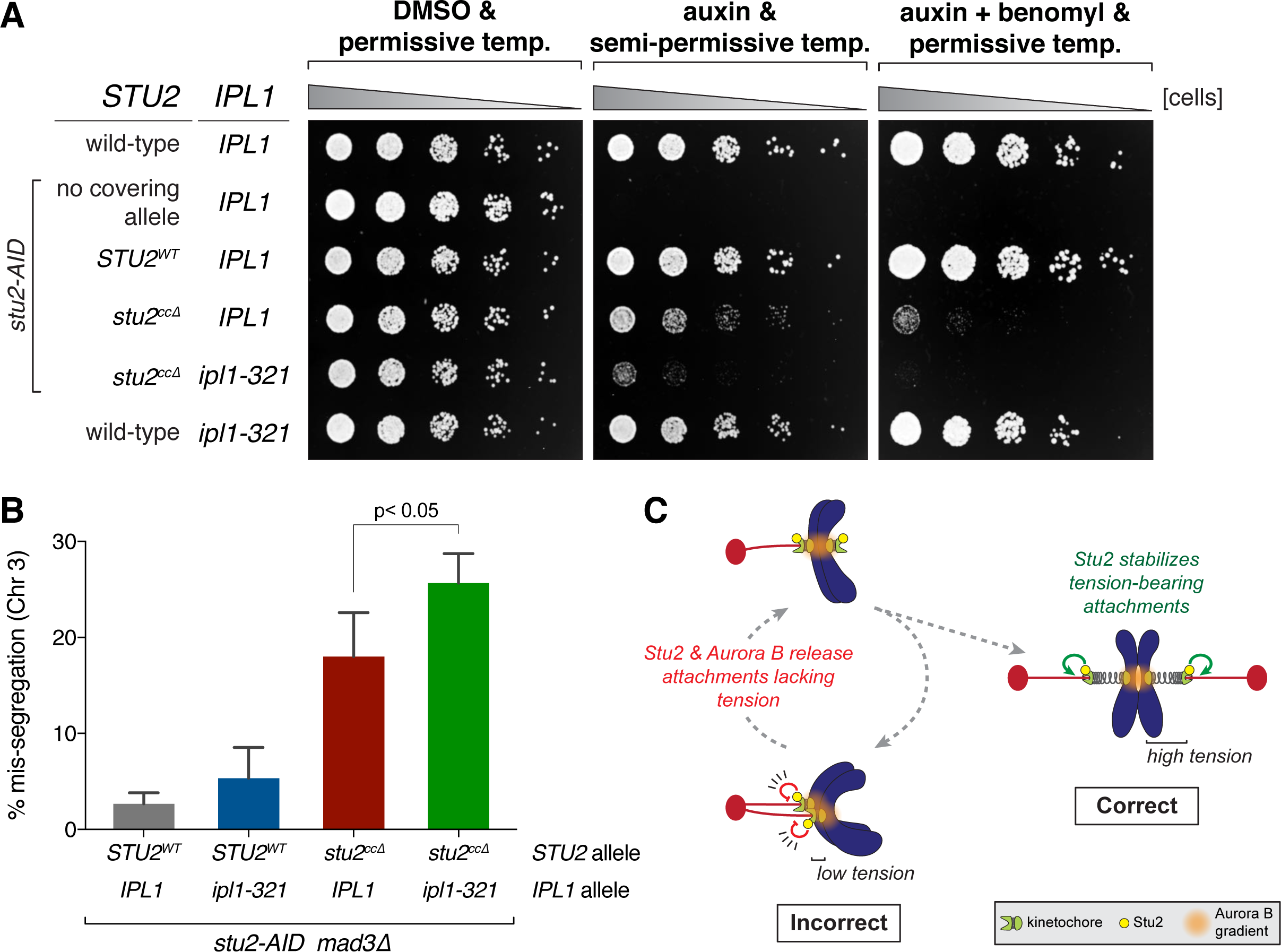
Mutants in Stu2 and Aurora B show a synergistic biorientation defect. A) Wild-type (SBY3), *stu2-AID* (no covering allele, SBY13772) and *stu2-AID* cells expressing various *STU2-3V5* alleles from an ectopic locus [*STU2^WT^*, SBY13903; *stu2^ccΔ^*, SBY13918] or also containing an *ipl1-321* allele (*stu2^ccΔ^ ipl1-321*, SBY17100) or *ipl1-321* alone (SBY630) were serially diluted (5-fold) and spotted on YPD or 5 µg/ml benomyl plates containing either DMSO or 500 µM auxin and incubated at 23°C (permissive) or 30°C (semi-permissive). B) Exponentially growing *stu2-AID mad3Δ* cells containing or lacking an *ipl1-321* allele and also an ectopically expressed *STU2* allele (*STU2^WT^ IPL1*, SBY18242; *STU2^WT^ ipl1-321*, SBY18244; *stu2^ccΔ^ IPL1*, SBY18246; *stu2^ccΔ^ ipl1-321*, SBY18248) that also contained a fluorescently labeled *CEN3* were arrested in G1 at the permissive temperature (23°C) and subsequently released from the G1 arrest into auxin containing media at a semi-permissive temperature (30°C). Quantification of chromosome mis-segregation in anaphase is shown. Error bars represent SD of three independent experiments; n = 100 cells for each experiment. p value was determined using a two-tailed paired t test. C) Model: Kinetochore-associated Stu2 confers tension sensitivity to kinetochore-microtubule attachments and is required for the establishment of bioriented attachments in vivo. Stu2 and Aurora B function together to release (destabilize) low tension bearing/incorrect kinetochore-microtubule attachments in cells. At high tension, a second function of Stu2 serves to stabilize correct attachments.

Finally, we asked whether the additive growth defects are correlated with an increase in chromosome mis-segregation. As above, we used the *stu2^ccΔ^* mutant and the *ipl1-321* allele to perturb these two pathways, and cells containing a fluorescently marked centromere of chromosome III to monitor chromosome segregation. We released cells from a G1 arrest into auxin containing media (to degrade the endogenous Stu2-AID protein) and at the semi-permissive temperature of 30°C and examined chromosome mis-segregation in anaphase (i.e. the percentage of cells that failed to segregate a copy of chromosome III to each of the resulting daughter nuclei). As expected, control cells expressing wild-type *STU2* and *IPL1*, as well as *ipl1-321* cells with the Aurora B pathway partially disrupted alone, displayed low levels of chromosome mis-segregation [*STU2^WT^ IPL1*, 2.7 ± 1.2%; *STU2^WT^ ipl1-321*, 5.3 ± 3.2%; Figure 6B]. Cells expressing the kinetochore localization-deficient *stu2^ccΔ^* mutant alone showed an enhanced level of chromosome missegregation [*stu2^ccΔ^ IPL1*, 18.0 ± 4.6%], however, the *stu2^ccΔ^ ipl1-321* double mutant cells displayed a significant increase in the rate of chromosome mis-segregation compared to either single mutant alone [*stu2^ccΔ^ ipl1-321*, 25.7 ± 3.1%; Figure 6B]. All together, these results suggest that the kinetochore’s intrinsic, Stu2-dependent, tension sensing pathway works in parallel with that of the canonical Aurora B-mediated error correction pathway. Furthermore, our findings are consistent with these two pathways both playing important roles in establishing correct bioriented attachments in vivo.

## DISCUSSION

The accurate segregation of chromosomes is a fundamental aspect of cell division. Ensuring that each daughter cell inherits the correct genomic complement requires that each pair of duplicated chromosomes attaches to microtubules from opposite poles of the cell, which will result in their proper disjunction during anaphase. The cell’s error correction machinery is responsible for promoting these bioriented attachments by ensuring that attachments under tension are stabilized. Here, we examine the extent to which the kinetochore’s intrinsic tension-sensing pathway, initially described in vitro (Akiyoshi et al., 2010; Miller et al., 2016), regulates kinetochore-microtubule attachments in vivo. We identify a mutant that abolishes kinetochore-associated Stu2 function, which we previously found was required for the intrinsic selectivity of tension-bearing kinetochore attachments (Miller et al., 2016). Disrupting these activities in vivo leads to error correction defects, indicating that the kinetochore’s intrinsic tension-sensing mechanism is important for establishing bioriented attachments in cells.

### Stu2 utilizes multiple domains to achieve kinetochore association

We previously found that, in vitro, the kinetochore’s intrinsic mechanical tension-sensing pathway requires dichotomous Stu2-dependent activities: promoting attachments to assembling microtubules when tension is high, and triggering detachment during microtubule disassembly at low force (Miller et al., 2016). To begin to understand how Stu2 carries out these complex functions, and the extent to which this intrinsic mechanism is participating in error correction in vivo, we determined the domain(s) within Stu2 that are required for kinetochore-association. The fact that Stu2 binds to both the kinetochore’s Ndc80 complex as well as microtubule plus ends has made this type of analysis challenging given their proximity in cells (Aravamudhan et al., 2014; Humphrey et al., 2018). Additionally, Stu2’s multiple functions in the cell has complicated our ability to examine its role in kinetochore function in cells since *stu2* loss of function mutants abolish spindle structure (Al-Bassam et al., 2006; Kosco et al., 2001; Pearson et al., 2003). Our use of kinetochore purifications and in vitro binding assays, in combination with cell biological assays examining mitotic spindle structure, allowed us to examine *stu2* mutants that are defective in kinetochore association but that maintain normal spindle length.

We identified two domains in Stu2 required for its kinetochore association, the extreme C-terminus and the coiled-coil domain, consistent with prior work (Haase et al., 2018; Humphrey et al., 2018). We propose that Stu2’s C-terminal tail domain (residues 855-888) directly interacts with Ndc80c, and that two copies of Stu2’s C-terminal domain are required to “sandwich” its binding motif on Ndc80c. Consistent with this idea, we observed numerous crosslinks between this domain and the Ndc80c (Figure 1C), and prior FRET-based observations also pointed to proximity between the C-terminus of Stu2 and the C-termini of Ndc80 and Nuf2 (Aravamudhan et al., 2014). However, the fact that the C-terminal tail domain has multiple binding partners that affect microtubule function (i.e. Bik1 and Bim1; (Stangier et al., 2018; Usui et al., 2003; Wolyniak et al., 2006) precluded further analysis of this mutant. How so many proteins compete for a small region of Stu2 raises interesting questions about cellular regulation that will need to be resolved in future studies. We also found that Stu2’s coiled-coil domain is also required for its kinetochore association, consistent with recent work in cells (Haase et al., 2018; Humphrey et al., 2018). Our observation that the bZIP allele restores kinetochore binding and viability suggest that it is homo-dimerization of Stu2 and not sequence specific elements within the coiled-coil domain that is important for Stu2’s kinetochore functions. While previous studies showed that Stu2’s in vitro microtubule binding is reduced when dimerization is prevented (Al-Bassam et al., 2006; Geyer et al., 2018), we found that the *stu2^ccΔ^* mutant cells maintained normal spindle length in vivo. Although we cannot rule out the possibility that the *stu2^ccΔ^* mutant has an effect on microtubule dynamics in vivo, the spindles appeared to function normally because inactivation of the spindle checkpoint allowed the *stu2^ccΔ^* mutant cells to enter anaphase and segregate chromosomes. Together, these results suggest that cells can either compensate for decreased Stu2 microtubule polymerase activity or that the polymerase activity of the dimerization-deficient mutant is normal in vivo (perhaps due to concentration differences in vivo and in vitro). Regardless, our finding that the *stu2^ccΔ^* mutant can support a normal spindle, yet is defective in kinetochore association, gave us a tool to examine Stu2’s kinetochore function in cells.

A critical still unanswered question is mechanistically how does Stu2 carry out its kinetochore function. Understanding the precise nature of the Stu2-Ndc80c interaction may begin to shed light on this question. For example, the kinetochore-bound pool of Stu2 may be held in a specific orientation that alters the manner in which it interacts with the microtubule tip, or perhaps affects the microtubule binding capacity of another component of the kinetochore. Determining the exact nature of this interaction, including the identity of the Ndc80 binding motif, will be an important question to address. We also observed that other regions of Stu2 may potentially be implicated in regulating its kinetochore function. Of particular interest, Stu2’s SK-rich region is completely dispensable for Stu2-kinetochore binding yet displays numerous crosslinks near Ndc80’s conserved hairpin region (Figure 1B-C). Recently, the SK-rich region was found to be involved in an intra-molecular regulation of Stu2, in which tubulin occupancy of the TOG domains regulates the ability of the SK-rich domain to bind to the microtubule lattice (Geyer et al., 2018). We also observed that while the N-terminus (i.e. TOG1) is not strictly required for Stu2’s ability to bind the kinetochore, we routinely observed decreased Stu2-kinetochore association in its absence (Figure 1B & S1C). An intriguing idea is that the intra-molecular regulation of Stu2 is involved in modulating Stu2’s kinetochore function. Perhaps intra-kinetochore tension and/or TOG domain-tubulin occupancy alters the ability of the SK-rich domain to interact with Ndc80’s hairpin domain, thus regulating the combined function of these proteins in microtubule tip binding. Future work examining mutations that perturb these domains will be required to determine whether these regions are involved in tuning Stu2’s kinetochore functions and the intrinsic tension-sensing pathway that it mediates.

### Multiple pathways promote biorientation in cells

Accurate chromosome segregation depends on the establishment of bioriented attachments. This process requires erroneous microtubule attachments to be selectively destabilized by the cell’s error correction machinery until all attachments become correctly bioriented. Error correction requires the activity of the conserved kinase Aurora B, but whether this pathway is solely responsible for correcting erroneous attachments has been less clear. We previously found that the kinetochore harbors an intrinsic tension-sensing mechanism such that high tension-bearing attachments are selectively stabilized, and this pathway depends on the conserved XMAP215 family member Stu2 (Akiyoshi et al., 2010; Miller et al., 2016). Whether this intrinsic tension-sensing mechanism regulates kinetochore-microtubule attachments in vivo, however, had yet to be determined. We have identified a Stu2 mutant that disrupts its kinetochore-association and kinetochore functions in vitro. Furthermore, we found that disrupting these activities in cells leads to error correction defects, and reminiscent of the requirements of Aurora B (Tanaka et al., 2002), this intrinsic tension-sensing pathway appears to be required for the establishment of bioriented attachments but dispensable for their maintenance. Finally, we found that this intrinsic mechanism works together with the Aurora B-mediated pathway in the process of error correction (Figure 6C). In the future, it will be critical to further investigate the relationship between these two error correction pathways, whether they function independently or temporally to promote biorientation and whether there is feedback between the two pathways. For example, Aurora B activity could regulate kinetochore-associated Stu2 function to tune the intrinsic tension-sensing pathway. Together, our findings suggest that the kinetochore’s intrinsic tension-sensing mechanism plays an important role in cells and that this pathway works together with the Aurora B-mediated mechanism to establish correctly bioriented attachments.

We propose that the kinetochore’s intrinsic tension-sensing mechanism constitutes a primordial error correction pathway in cells. Kinetochore-associated Stu2 enables long-lived kinetochore attachments when tension is high and ensures that non-tension bearing kinetochore attachments are short-lived. These combined activities result in an error correction mechanism that promotes bioriented attachments, and perhaps primitive cells relied on this pathway to ensure accurate chromosome segregation prior to evolving the Aurora B-mediated pathway.

## Conclusions

Our findings suggest that the kinetochore’s intrinsic tension-sensing mechanism, in which high tension bearing attachments are directly stabilized, plays an important role in cells and works with the canonical Aurora B-mediated error correction pathway to promote accurate chromosome segregation. Given that XMAP215 family members, including human ch-TOG, display a conserved association with the kinetochore’s Ndc80c (Aravamudhan et al., 2014; Haase et al., 2018; Hsu and Toda, 2011; Humphrey et al., 2018; Miller et al., 2016; Tang et al., 2013), it will be important to investigate whether intrinsic tension-selectivity is a wide-spread feature of kinetochore-microtubule interactions, whether mutations that alter ch-TOG regulation (Charrasse et al., 1995, 1998) lead to defects in error correction, and whether these defects correlate with the increased rates of chromosome mis-segregation observed in most cancers.

## ACKNOWLEDGEMENTS

We are grateful to Arshad Desai for providing antibodies, Angelika Amon, Leon Chan, Sue Jaspersen, Frank Uhlmann, Eris Duro, and Adèle Marston for reagents, and to Ajit Joglekar for helpful discussions and sharing data prior to publication. M.P.M. is an HHMI Fellow of the Damon Runyon Cancer Research Foundation. This work was supported by NIH grants T32CA080416 (to M.P.M.), P41GM103533 (to M.J.M.), R01GM098543 (to L.M.R.), R01GM040506 (to T.N.D.), R01GM079373 and P01GM105537 (to C.L.A.), and R01GM064386 (to S.B.). Also, this work was supported by the Genomics and Scientific Imaging Shared Resources of the Fred Hutch/University of Washington Cancer Consortium (P30 CA015704). E.A.G. was supported by T32GM008297 from NIH, and by an NSF Graduate Research Fellowship, Grant No. 2014177758. C.L.A. is supported by a David and Lucile Packard Fellowship (2006-30521). S.B. is also an investigator of the Howard Hughes Medical Institute.

## FIGURE LEGENDS

**Supplement to Figure 1.**
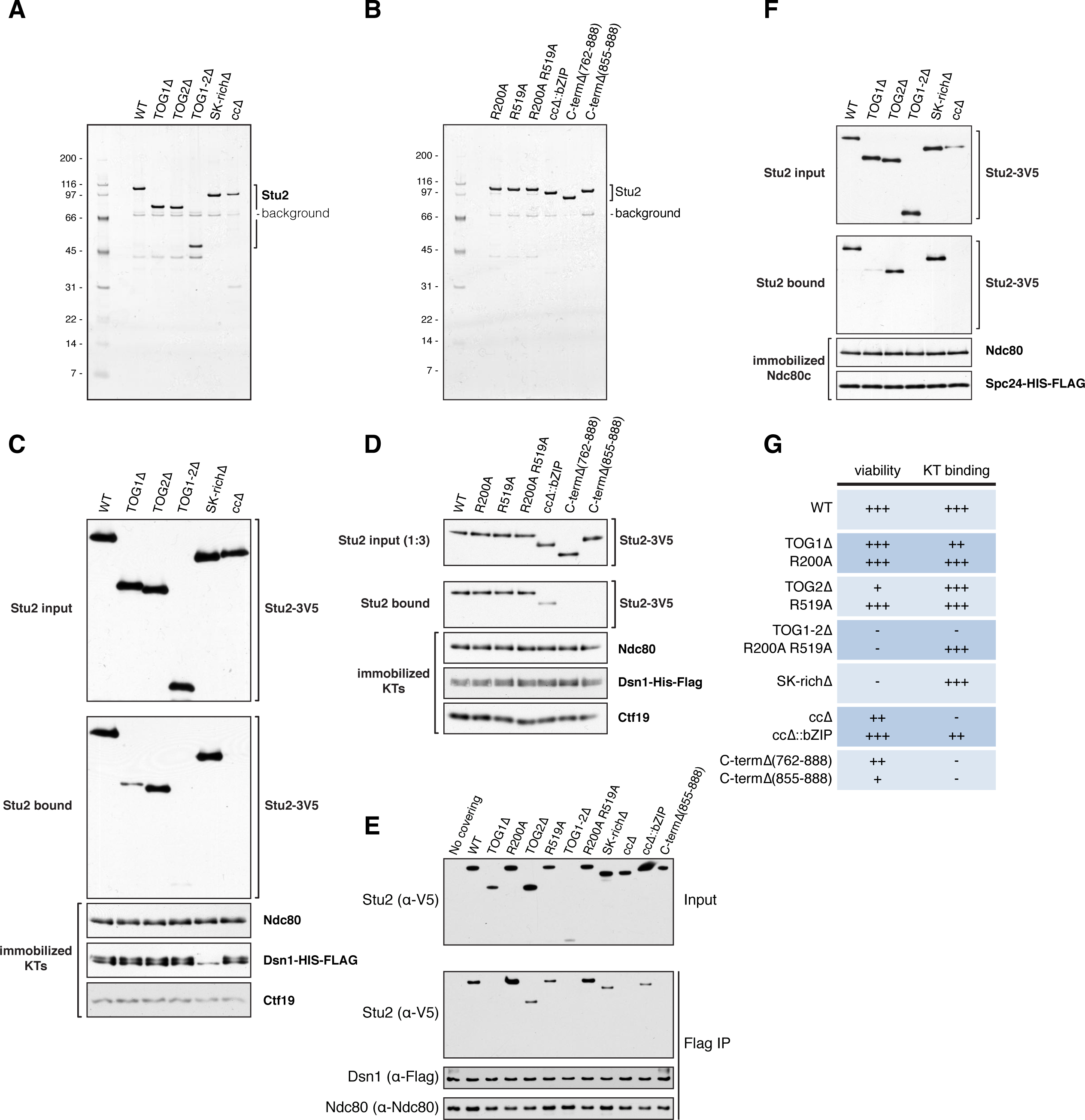
Stu2’s kinetochore association depends on homo-dimerization and its extreme C-terminus. A) Protein lysates were prepared from exponentially growing *stu2-AID* cultures expressing various *STU2-3V5* alleles from an ectopic locus treated with auxin 30 min prior to harvesting (*STU2^WT^*, SBY13557; *stu2^TOG1Δ^*, SBY13563; *stu2^TOG2Δ^*, SBY13569; *stu2^TOG1-2Δ^*, SBY13575; *stu2^SK-richΔ^*, SBY13581; *stu2^ccΔ^*, SBY13587). Stu2-3V5 was purified by α-V5 IP, followed by washes in buffer containing 1.0 M KCl (BH 1.0) then V5 peptide elution. Eluate was run on an SDS-PAGE gel and analyzed by silver stain. Background bands were previously determined by mass spectrometry to be the highly homologous heat shock proteins Ssa1, Ssa2 (70 kDa), and Ssb1, Ssb2 (66 kDa), which are common co-purifying proteins in IPs from yeast lysates (Cheeseman et al., 2002), and were isolated in all purifications. B) As in (A) expressing different *STU2-3V5* alleles (*stu2^R200A^*, SBY13923; *stu2^R519A^*, SBY13929; *stu2^R200A R519A^*, SBY13933; *stu2^ccΔ::bZIP^*, SBY13939; *stu2^C-termΔ(762-888)^*, SBY14267; *stu2^C-termΔ(855-888)^*, SBY14273). C) Protein lysates were prepared from exponentially growing cultures containing Dsn1-6His-3Flag Stu2-AID (SBY13772) treated with 500 μM auxin for 30 min prior to harvesting cells. Kinetochore particles were immobilized by α-Flag IP. Immobilized kinetochore-beads were incubated with 30 ng of Stu2-3V5 variants (purified as in A) for 30 min at room temperature, washed and eluted with Flag peptide. Kinetochore-bound proteins were analyzed by immunoblotting with α-Flag, α-V5, α-Ndc80 and α-Ctf19 antibodies. Note: Stu2^SK-richΔ^ and Dsn1-6His-3Flag co-migrate on an SDS-PAGE gel. For technical reasons (that are not entirely clear), detection of Dsn1-6His-3Flag was affected by first probing for Stu2^SK-richΔ^, however similar Dsn1-6His-3Flag levels were observed for all samples when these sample eluates were run on an SDS-PAGE gel and analyzed by silver stain. D) As in (C) using Stu2-3V5 variants (purified as in B). E) Exponentially growing *stu2-AID* cultures expressing an ectopic copy of Stu2 (*STU2^WT^*, SBY13901; *stu2^TOG1Δ^*, SBY13904; *stu2^R200A^*, SBY13919; *stu2^TOG2Δ^*, SBY13907; *stu2^R519A^*, SBY13925; *stu2^TOG1-2Δ^*, SBY13910; *stu2^R200A R519A^*, SBY13930; *stu2^SK-richΔ^*, SBY13913; *stu2^ccΔ^*, SBY13916; *stu2^ccΔ::bZIP^*, SBY13935 or *stu2^C-termΔ(855-888)^*, SBY14269) or without an ectopic allele (no covering allele, SBY13772) that also contained Dsn1-6His-3Flag were treated with auxin 30 min prior to harvesting. Protein lysates were subsequently prepared and kinetochore particles were purified by α-Flag immunoprecipitation (IP) and analyzed by immunoblotting. F) Protein lysates were prepared from exponentially growing cultures containing Spc24-6His-3Flag Spc105-AID (SBY14022) treated with 500 μM auxin for 30 min prior to harvesting cells. Ndc80c was immobilized by α-Flag IP. Immobilized Ndc80c-beads were incubated with 30 ng of Stu2-3V5 variants (purified as in A) for 30 min at room temperature, washed and eluted with Flag peptide. Ndc80c-bound proteins were analyzed by immunoblotting with α-Flag, α-V5, and α-Ndc80 antibodies. G) Summary table of kinetochore binding and viability data from Figures 1, 2, S1 and S2.

**Supplement to Figure 2.**
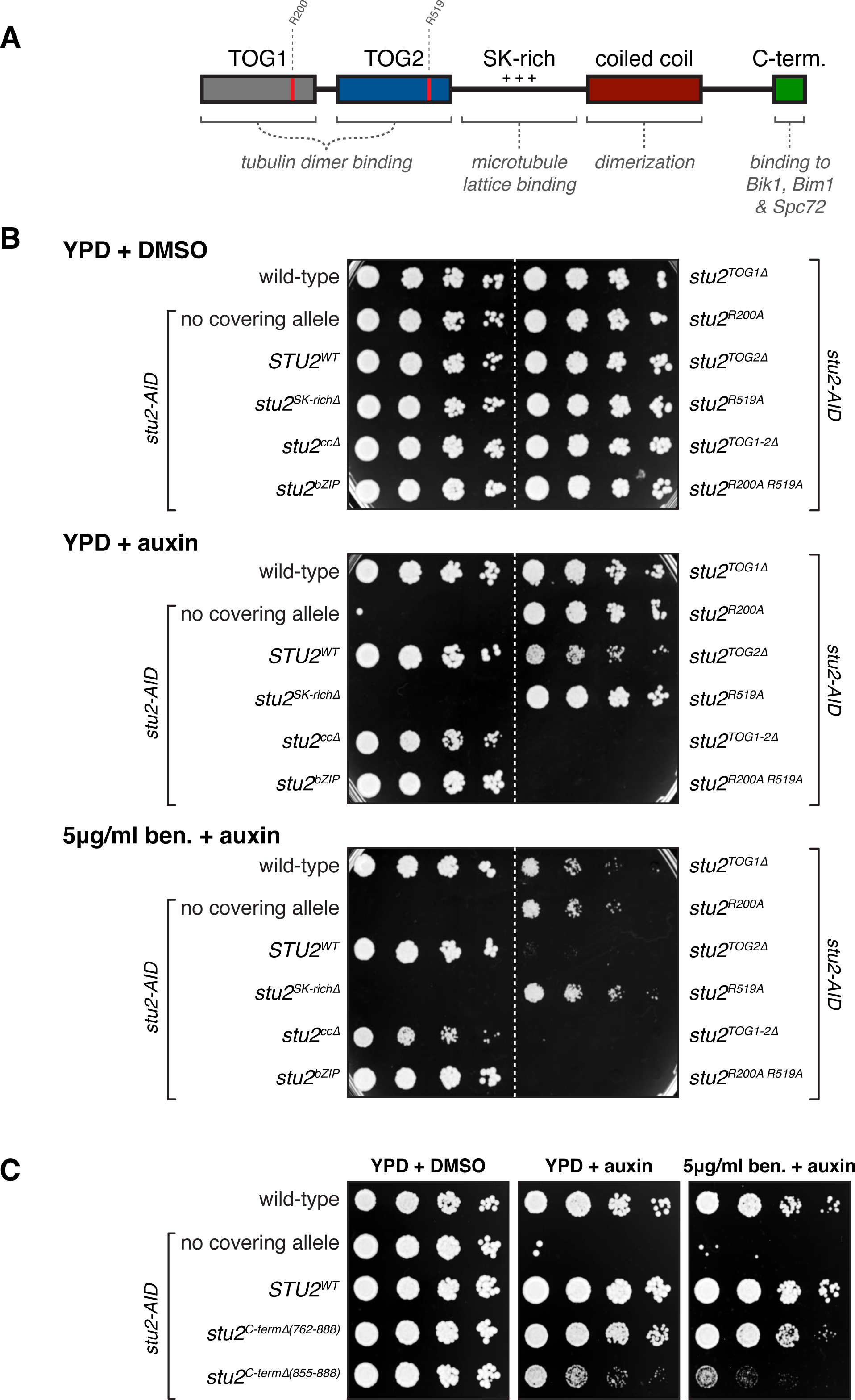
Phenotype of various *stu2* mutants. A) Schematic of Stu2’s domain architecture. B) Wild-type (SBY3), *stu2-AID* (no covering allele, SBY13772) and *stu2-AID* cells expressing various *STU2-3V5* alleles from an ectopic locus (*STU2^WT^*, SBY13901; *stu2^SK-richΔ^*, SBY13913; *stu2^ccΔ^*, SBY13916; *stu2^ccΔ::bZIP^*, SBY13935; *stu2^TOG1Δ^*, SBY13904; *stu2^R200A^*, SBY13919; *stu2^TOG2Δ^*, SBY13907; *stu2^R519A^*, SBY13925; *stu2^TOG1-2Δ^*, SBY13910; *stu2^R200A R519A^*, SBY13930) were serially diluted (5-fold) and spotted on YPD or 5 µg/ml benomyl plates containing either DMSO or 100 µM auxin. C) Wild-type (SBY3), *stu2-AID* (no covering allele, SBY13772) and *stu2-AID* cells expressing various *STU2-3V5* alleles from an ectopic locus [*STU2^WT^*, SBY13901; *stu2^C-termΔ(762-888)^*, SBY14263; *stu2^C-termΔ(855-888)^*, SBY14269) were serially diluted (5-fold) and spotted on YPD or 5 µg/ml benomyl plates containing either DMSO or 100 µM auxin.

**Supplement to Figure 4.**
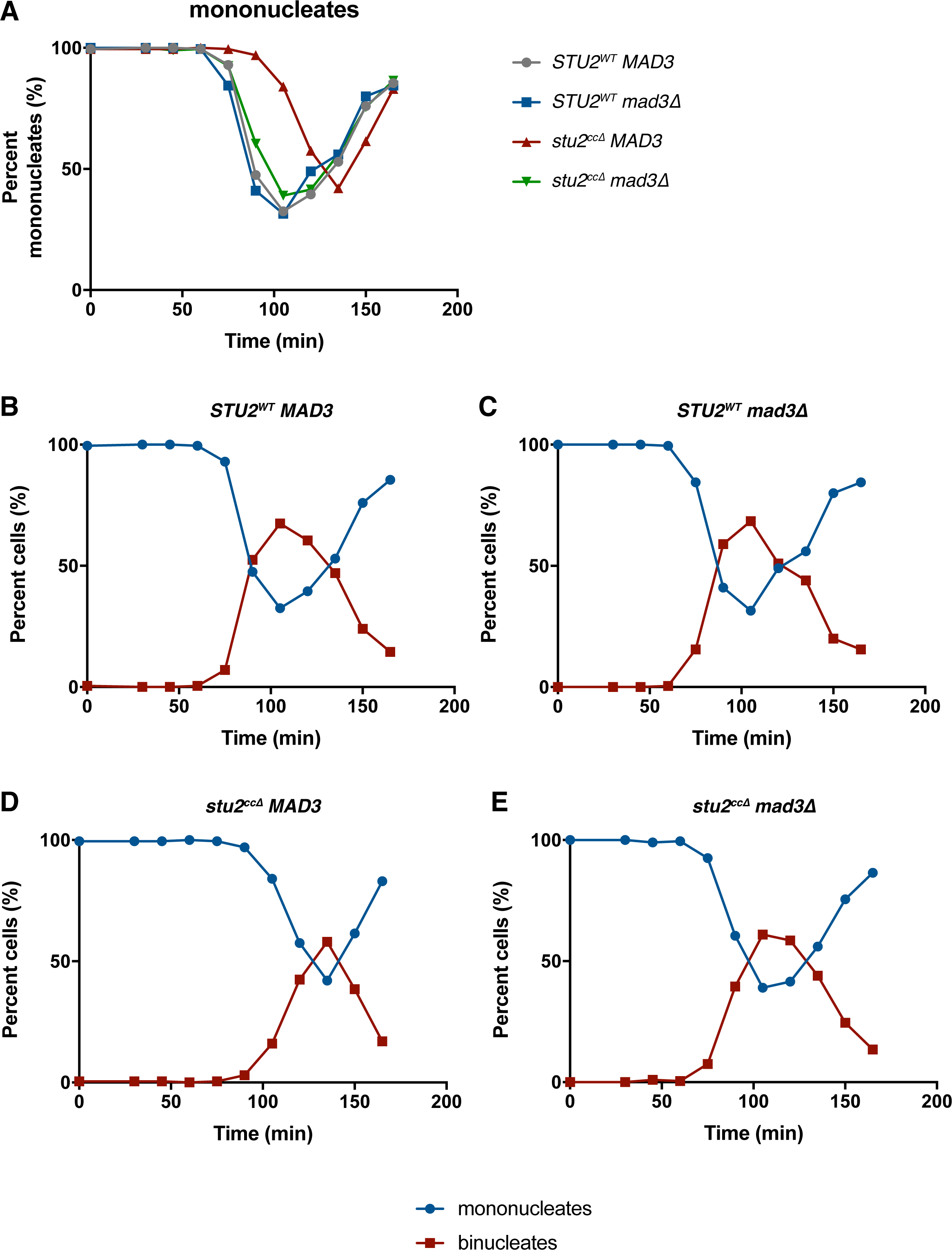
Cell cycle progression of kinetochore-binding deficient Stu2 mutant. A-E) Cell cycle progression from Figure 4D. Quantification of the number of mononucleate and binucleate cells for each strain. Shown is a representative experiment.

**Supplement to Figure 5.**
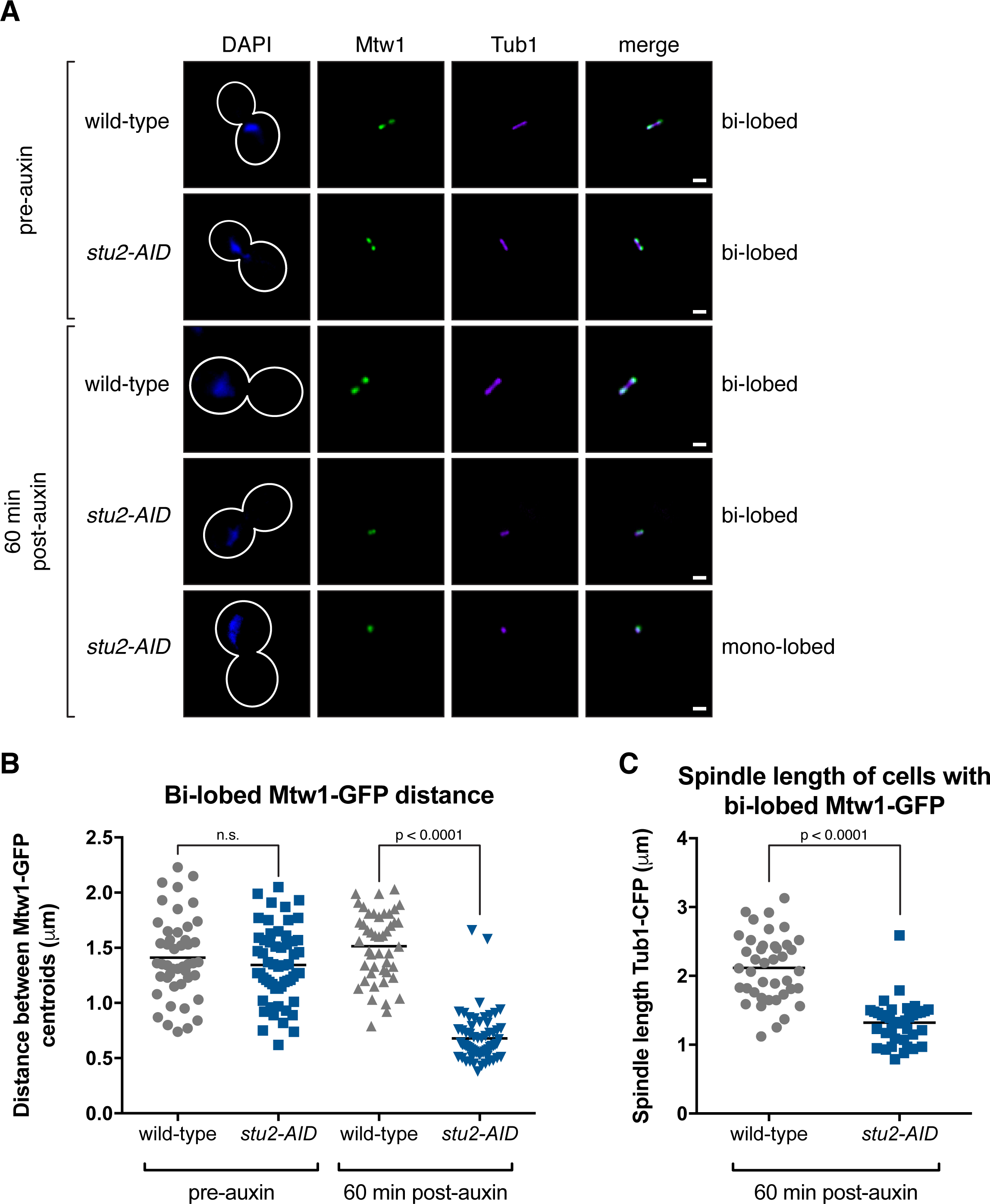
Stu2 function is required to maintain a normal bipolar spindle and bi-lobed kinetochore distribution. We used the Stu2-AID system to examine the effect of depleting all cellular Stu2 on spindle structure after the cells had already formed a mitotic spindle. For these experiments, we arrested cells in metaphase by depleting Cdc20 (now using a methionine repressible *pMET-CDC20* allele) and found that the subsequent degradation of the Stu2-AID protein led to a significant decrease in both spindle length and collapse of the bi-lobed kinetochore clusters to a mono-lobed focus (Figure S5A-C). A recent study used an “anchor away” system to address this same question (Humphrey et al., 2018). However, we repeated this experiment because that study only observed a 70% mis-localization of Stu2 by fluorescence microscopy and no alteration in mitotic spindle length, suggesting incomplete depletion of Stu2 from the nucleus. A) Exponentially growing cells carrying *pMET-CDC20 MTW1-3GFP TUB1-CFP* and either the wild-type *STU2* (SBY17105) or *stu2-AID* allele (SBY17106) at the endogenous *STU2* locus were arrested in metaphase by depleting Cdc20 (by the addition of methionine to the media). Once cells were arrested in metaphase, auxin was added to induce degradation of the Stu2-AID protein, and the cells were subsequently analyzed for kinetochore distribution and spindle morphology (by examining Mtw1-3GFP and Tub1-CFP, respectively). Representative images for each are shown for pre-auxin and 60 min post-auxin addition. Kinetochore distribution was determined to be bi- or mono-lobed. B) Kinetochore distribution (distance between bi-lobed kinetochore clusters) was measured for cells described in (A). n = 46-63 cells; p values were determined using a two-tailed unpaired t test (n.s. = not significant). C) Spindle length (length of Tub1-CFP) was measured for cells described in (A). n = 38-41 cells; p value was determined using a two-tailed unpaired t test.

**Supplement to Figure 6.**
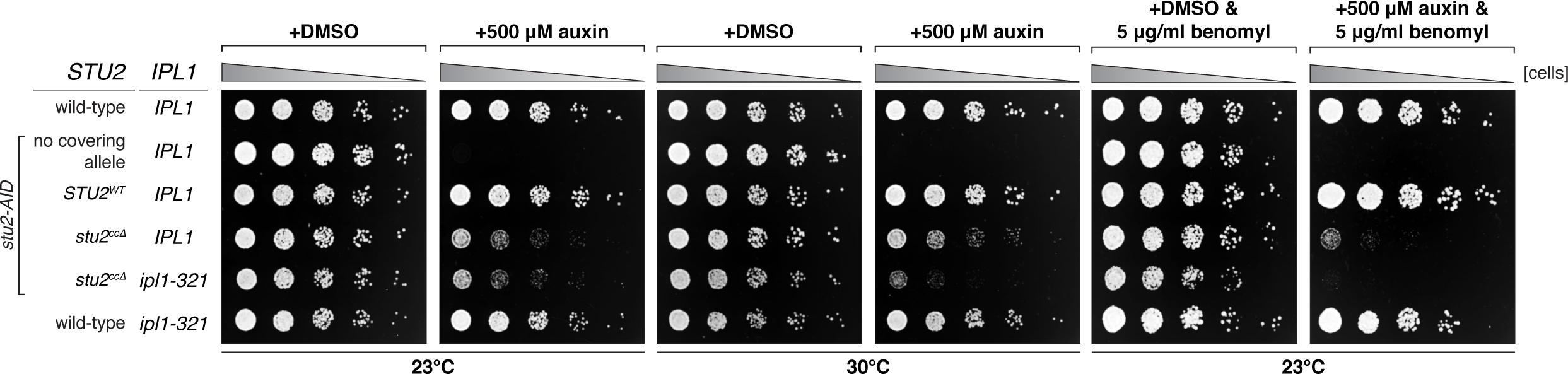
*stu2^ccΔ^* mutant displays synthetic phenotype with an Aurora B mutant. Wild-type (SBY3), *stu2-AID* (no covering allele, SBY13772) and *stu2-AID* cells expressing various *STU2-3V5* alleles from an ectopic locus (*STU2^WT^*, SBY13903; *stu2^ccΔ^*, SBY13918) or also containing an *ipl1-321* allele (*stu2^ccΔ^ ipl1-321*, SBY17100) or *ipl1-321* alone (SBY630) were serially diluted (5-fold) and spotted on YPD or 5 µg/ml benomyl plates containing either DMSO or 500 µM auxin and incubated at 23°C (permissive) or 30°C (semi-permissive).

## MATERIALS AND METHODS

### Strain Construction and Microbial Techniques

#### Yeast Strains and Plasmids

*Saccharomyces cerevisiae* strains used in this study are described in Table S1 and are derivatives of SBY3 (W303). Standard media and microbial techniques were used (Sherman et al., 1974). Yeast strains were constructed by standard genetic techniques. Construction of *pCUP1-GFP-LacI* and *ipl1-321* are described in (Biggins et al., 1999), *CEN3::lacO:TRP1* and *CEN8::lacO:TRP1* are described in (Shonn et al., 2003), *MTW1-3GFP, TUB1-CFP* and *mad3Δ* are described in (Pinsky et al., 2006), *DSN1-6His-3Flag* is described in (Akiyoshi et al., 2010), *stu2-3V5-IAA7*, *stu2-3HA-IAA7, and SPC24-6His-3Flag* were constructed by PCR-based methods (Longtine et al., 1998) and are described in (Miller et al., 2016). Strains containing previously described alleles were also generously provided (*cdc20-IAA17* from Eris Duro and Adèle Marston; *SPC42-CFP* from Sue Jaspersen; *pMET-CDC20* from Frank Uhlmann). *pADH1-TIR1-9Myc* is described in (Nishimura et al., 2009). *pGPD1-TIR1* integration plasmids (pSB2271 for integration at *LEU2*, pSB2273 for integration at *HIS3* or pSB2275 for integration at *TRP1*) as well as a 3V5-IAA7 tagging plasmid (pSB2065) were provided by Leon Chan. Construction of a 3HA-IAA7 tagging plasmid (pSB2229) was described previously in (Miller et al., 2016). Construction of a *LEU2* integrating plasmid containing wild-type *pSTU2-STU2-3V5* (pSB2232) is also described in (Miller et al., 2016). *STU2* variants were constructed by mutagenizing pSB2232 as described in (Liu and Naismith, 2008; Tseng et al., 2008), resulting in pSB2254 (*pSTU2-stu2(Δ2-281)-3V5*, i.e. *stu2^TOG1Δ^*), pSB2255 (*pSTU2-stu2(Δ2-550)-3V5*, i.e. *stu2^TOG1-2Δ^*), pSB2257 (*pSTU2-stu2(Δ282-550::GDGAGL^linker^)-3V5*, i.e. *stu2^TOG2Δ^*), pSB2259 (*pSTU2-stu2(Δ551-657::GDGAGL^linker^)-3V5*, i.e. *stu2^SK-richΔ^*), pSB2261 (*pSTU2-stu2(Δ658-761::GDGAGL^linker^)-3V5*, i.e. *stu2^ccΔ^*), pSB2306 (*pSTU2-stu2(R200A)-3V5*, i.e. *stu2^R200A^*), pSB2307 (*pSTU2-stu2(R519A)-3V5*, i.e. *stu2^R519A^*), pSB2308 (*pSTU2-stu2(R200A R519A)-3V5,* i.e. *stu2^R200A R519A^*), pSB2309 (*pSTU2-stu2(Δ658-761::GDGAGL^linker^-GCN4(249-281)^bZIP^-GDGAGL^linker^)-3V5*, i.e. *stu2^ccΔ::bZIP^*), pSB2357 (*pSTU2-stu2(Δ762-888)-3V5*, i.e. *stu2^C-termΔ(762-888)^*), pSB2358 (*pSTU2-stu2(Δ855-888)-3V5*, i.e. *stu2^C-termΔ(855-888)^*). Further details of plasmid construction including plasmid maps and primer sequences available upon request.

#### Auxin Inducible Degradation

The <u>a</u>uxin inducible <u>d</u>egron (AID) system was used essentially as described (Nishimura et al., 2009). Briefly, cells expressed C-terminal fusions of the protein of interest to an auxin responsive protein (IAA7 or IAA17) at the endogenous locus. Cells also expressed TIR1, which is required for auxin-induced degradation. 100-500 μM IAA (indole-3-acetic acid dissolved in DMSO; Sigma) was added to media to induce degradation of the AID-tagged protein. Auxin was added for 30 min prior to harvesting cells or as is indicated in figure legends.

#### Spotting Assay

For the spotting assay, the desired strains were grown overnight in yeast extract peptone plus 2% glucose (YPD) medium. The following day, cells were diluted to OD_600_ ~1.0 from which a serial 1:5 dilution series was made and spotted on YPD+DMSO or YPD+100-500 μM IAA (indole-3-acetic acid dissolved in DMSO) plates. Plates were incubated at 23-30°C for 2-3 days unless otherwise noted.

#### Cell Fixation and Imaging Conditions

Exponentially growing cultures were treated with 500 μM auxin for 2h, then fixed (see below) and analyzed for Mtw1-3GFP localization and spindle morphology (Spc42-CFP or Tub1-CFP). An aliquot of cells was fixed with 3.7% formaldehyde in 100mM phosphate buffer (pH 6.4) for 5 min. Cells were washed once with 100mM phosphate (pH 6.4), resuspended in 100mM phosphate, 1.2M sorbitol buffer (pH 7.5) and permeabilized with 1% Triton X-100 stained with 1 μg/ml DAPI (4′, 6-diamidino-2-phenylindole; Molecular Probes). Cells were imaged using a Nikon E600 microscope with a 60X objective (NA=1.40), equipped with a Photometrics Cascade 512B digital camera. Five Z-stacks (0.3 micron apart) were acquired and all frames with nuclear signal in focus were maximally projected. NIS Elements software (Nikon) was used for image acquisition and processing.

#### Biorientation Assay

In metaphase, sister kinetochores become bioriented and are transiently stretched apart by opposing microtubule pulling forces (Goshima and Yanagida, 2000; He et al., 2001; Pearson et al., 2001), which can be visualized by fluorescently marking the centromere of a single chromosome (Straight et al., 1996). Biorientation was examined in metaphase arrested cells as judged by the fluorescently marked centromeres appearing as two distinct foci. Cells were grown in YPD (for *cdc20-AID* containing strains) or media lacking methionine (for *pMET-CDC20* containing strains in Figure 5). Exponentially growing cells were subsequently arrested in metaphase either by the addition of 500μM IAA for 2h (for *cdc20-AID* containing strains in Figure 4C, whereby biorientation was examined); or the addition of 8mM methionine each hour for 3h (for *pMET-CDC20* containing strains in Figure 5). The *maintenance* of biorientation was determined in *pMET-CDC20* containing strains by the addition of 500μM IAA (to degrade endogenous Stu2-AID protein) for 2h and monitoring biorientation over time.

#### Cell Cycle Progression and Chromosome Segregation Assay

Cells were grown in YPD medium. Exponentially growing *MATa* cells also carrying a tandem array of lacO sequences integrated either proximal to *CEN3* or *CEN8* (Shonn et al., 2003) and a LacI-GFP fusion (Biggins et al., 1999; Straight et al., 1996) were arrested in G1 with 1μg/ml α-factor. When arrest was complete, cells were released into medium lacking α-factor pheromone and containing 500μM IAA at either 23°C or 30°C (semi-permissive temperature). ~75 min after G1 release (for 23°C) or ~65 min after G1 release (for 30°C), 1μg/ml α-factor was added to prevent a second cell division. Samples were taken every 15min after G1 release to determine cell cycle progression (via nuclear morphology of DAPI stained nuclei) and chromosome segregation in anaphase (for Figure 4E at 23°C, chromosome segregation at 120 min post-release is quantified; for Figure 6B at 30°C, chromosome segregation at 75 min post-release is quantified).

### Protein Biochemistry

#### Purification of Native Kinetochore Particles, Ndc80 Complex and Stu2

Native kinetochore particles, Ndc80c and Stu2 were purified from asynchronously growing *S. cerevisiae* cells as described below (unless otherwise noted in the text). For kinetochore particles, an α-Flag immunoprecipitation of Dsn1-6His-3Flag was performed (essentially as described in (Akiyoshi et al., 2010). To purify the Ndc80c, an α-Flag immunoprecipitation of Spc24-6His-3Flag was performed (protein lysate from SBY14022). To purify Stu2, an α-V5 immunoprecipitation of the Stu2-3V5 variant was performed from strains in which the endogenous Stu2-AID protein was degraded (see below). For each, cells were grown in yeast peptone dextrose rich (YPD) medium. For strains containing Stu2-AID or Spc105-AID, cells were treated with 500 μM auxin 30 min prior to harvesting. Protein lysates were prepared by lysing cells in a Freezer/Mill (SPEX SamplePrep) submerged in liquid nitrogen (Sarangapani et al., 2014) or mechanically disrupted in the presence of lysis buffer using glass beads and a beadbeater (Biospec Products). Lysed cells were resuspended in buffer H (BH) (25 mM HEPES pH 8.0, 2 mM MgCl_2_, 0.1 mM EDTA, 0.5 mM EGTA, 0.1% NP-40, 15% glycerol with 150 mM KCl for native kinetochores, 750 mM KCl for native Ndc80c or 1 M KCl for native Stu2) containing protease inhibitors (at 20 μg mL^−1^ final concentration for each of leupeptin, pepstatin A, chymostatin and 200 μM phenylmethylsulfonyl fluoride) and phosphatase inhibitors (0.1 mM Na-orthovanadate, 0.2 μM microcystin, 2 mM β-glycerophosphate, 1 mM Na pyrophosphate,5 mM NaF) followed by ultracentrifugation at 98,500 g for 90 min at 4 °C. Lysates prepared using the beadbeater were instead centrifuged at 16,100 g for 30 min at 4 °C. Dynabeads conjugated with α-Flag or α-V5 antibodies were incubated with extract for 3 h with constant rotation, followed by three washes with BH containing protease inhibitors, phosphatase inhibitors, 2 mM dithiothreitol (DTT) and either 150 mM KCl (kinetochores) or 1 M KCl (Ndc80c and Stu2). Beads were further washed twice with BH containing 150 mM KCl and protease inhibitors. Associated proteins were eluted from the beads by gentle agitation of beads in elution buffer (0.5 mg ml^−1^ 3Flag peptide or 0.5 mg ml^−1^ 3V5 peptide in BH with 150 mM KCl and protease inhibitors) for 30 min at room temperature. To eliminate the co-purification of the Spc105 complex with the native Ndc80c, we immunoprecipitated Spc24-6His-3Flag from cells carrying an *spc105-AID* allele that were treated with 500 μM auxin for 30 min.

#### Immunoblot Analysis

For immunoblot analysis, cell lysates were prepared as described above (Protein biochemistry section) or by pulverizing cells with glass beads in sodium dodecyl sulfate (SDS) buffer using a bead-beater (Biospec Products). Standard procedures for sodium dodecyl sulfate-polyacrylamide gel electrophoresis (SDS-PAGE) and immunoblotting were followed as described in (Burnette, 1981; Towbin et al., 1992). A nitrocellulose membrane (BioRad) was used to transfer proteins from polyacrylamide gels. Commercial antibodies used for immunoblotting were as follows: α-Flag, M2 (Sigma-Aldrich) 1:3,000; α-V5 (Invitrogen) 1:5,000. Antibodies to Ctf19, and Ndc80 were kind gifts from Arshad Desai and were used at: α-Ctf19, (OD10) 1:1,000; and α-Ndc80, (OD4) 1:10,000. The secondary antibodies used were a sheep anti-mouse antibody conjugated to horseradish peroxidase (HRP) (GE Biosciences) at a 1:10,000 dilution or a donkey anti-rabbit antibody conjugated to HRP (GE Biosciences) at a 1:10,000 dilution. Antibodies were detected using the SuperSignal West Dura Chemiluminescent Substrate (Thermo Scientific).

#### In vitro Binding Assays

To examine the binding of Stu2 to purified kinetochores, Stu2-3V5 and native kinetochores were purified from exponentially growing yeast cells as described above (Protein biochemistry section). Prior to eluting purified kinetochores (with approximately 75-100 ng of Dsn1-Flag bound) from α-Flag dynabeads, beads were incubated with 15 μl (30-90 ng) of purified Stu2-3V5 for 30 min at room temperature with gentile agitation. Beads were then washed twice with BH containing 150 mM KCl and protease inhibitors. Associated proteins were eluted from the beads by gentle agitation of beads in elution buffer (0.5 mg ml^−1^ 3Flag peptide in BH with 150 mM KCl and protease inhibitors) for 30 min at room temperature.

#### Recombinant Protein Expression and Purification

The *S. cerevisiae* Ndc80c was expressed in *E. coli* using polycistronic vectors, and purified as previously described (Powers et al., 2009; Tien et al., 2010; Wei et al., 2005). Stu2 constructs were made in pHAT vector containing N-terminal 6His tag, C-terminal eGFP-tag followed by a *Strep*-tag II (the original vector was a gift from Dr. Gary Brouhard). Expression was induced in *E. coli* using Arctic Express Cells and purified as in (Geyer et al., 2018).

#### Chemical Cross-linking and Mass Spectrometry Analysis (XL-MS)

Cross-linking reactions, mass spectrometry and data analysis were carried out as described previously (Zelter et al., 2015). A protein mixture containing 3.3 μg Stu2 plus 3 μg Ndc80c in a final volume of 44 μL BRB80 (80 mM PIPES, 1mM EGTA, 1 mM MgCl_2_, 10 μM taxol) was incubated for 15 minutes at room temperature. Cross-linking was initiated by adding 3.75 μL of 145 mM EDC plus 1.88 μL of 145 mM sulfo-NHS dissolved in BRB80. Cross-linking was allowed to proceed for 30 minutes at room temperature before quenching by addition of 5 μL 1M ammonium bicarbonate plus 0.5 μL 2-mercaptoethanol. Samples were reduced with 10 mM dithiothreitol (DTT) for 30 mins at 37°C followed by alkylation with 15 mM iodoacetamide (IAA) for 30 mins at room temperature. Tryptic digestion was performed at 37oC for 4 hours with shaking at a substrate to trypsin ratio of 18:1. After digestion, samples were acidified with 5 M HCl and stored at −80°C until analysis. Mass spectrometry was performed on a Q-Exactive HF (Thermo Fisher Scientific) in data dependent mode by loading 0.4 μg of sample onto a fused-silica capillary tip column (75-μm i.d.) packed with 30 cm of Reprosil-Pur C18-AQ (3-μm bead diameter, Dr. Maisch). Peptides were eluted from the column at 0.25 μl/min using a 120 min acetonitrile gradient. Spectra were converted into mzML using msconvert from ProteoWizard (Chambers et al., 2012). Proteins present in the sample were identified using Comet (Eng et al., 2013). Cross-linked peptides were identified within those proteins using Kojak version 1.4.3 (Hoopmann et al., 2015) available at (http://www.kojak-ms.org).

Percolator version 2.08 (Käll et al., 2007) was used to assign a statistically meaningful q value to each peptide spectrum match (PSM) through analysis of the target and decoy PSM distributions. The target databases consisted of all proteins identified in the sample. The decoy database consisted of the corresponding set of reversed protein sequences. Data presented here were filtered to show hits to the target proteins that had a Percolator assigned peptide level q value ≤ 0.05. The complete, unfiltered list of all PSMs and their Percolator assigned q values, are available on the ProXL web application (Riffle et al., 2016) at: https://proxl.yeastrc.org/proxl/viewProject.do?project_id=52 along with the raw MS spectra and search parameters used.

### Optical Trap Assays

#### Bead Preparation for Optical Trap Assays

Optical trap-based bead motility assays were performed as in (Akiyoshi et al., 2010; Miller et al., 2016; Umbreit et al., 2012). Streptavidin-coated 0.44-μm polystyrene beads (Spherotech) were functionalized with biotinylated anti-penta-His antibody (Qiagen) and decorated with purified native Ndc80c (via Spc24-6His-3Flag). Bead decoration was performed in a total volume of 20 μl incubation buffer (BRB80 containing 1 mg ml^−1^ κ-casein). 10 nM of purified native Ndc80c was incubated with 6 pM beads for 1 h at 4 °C, unbound protein was removed by pelleting the beads (16,000 g for 10 min at 4 °C), washing with ~200 μl of incubation buffer, pelleting beads again (16,000 g for 10 min at 4 °C) and resuspending in original volume. For the addition of purified Stu2 to Ndc80c, Ndc80c-decorated beads were prepared as above and purified Stu2-3V5 was added to the microtubule growth buffer (see below) to a final concentration of 2 nM as in (Miller et al., 2016).

#### Rupture Force Measurements

Dynamic microtubule extensions were grown from coverslip-anchored GMPCPP-stabilized microtubule seeds in a microtubule growth buffer consisting of BRB80, 1 mM GTP, 250 µg ml^−1^ glucose oxidase, 25 mM glucose, 30 µg ml^−1^ catalase, 1 mM DTT, 1.4-1.5 mg ml^−1^ purified bovine brain tubulin, and blocking protein (1 mg ml^−1^ κ-casein). Assays were performed at 23 °C. Rupture force experiments were performed as in (Akiyoshi et al., 2010; Miller et al., 2016; Sarangapani et al., 2013, 2014). Briefly, an optical trap was used to apply a force of ∼2 pN in the direction of microtubule assembly. Once beads were observed to track with microtubule growth for a distance of ~100-300 nm (to ensure end-on attachment), the applied force was increased at a constant rate of 0.25 pN s^−1^ until bead detachment. Records of bead position over time were collected and analyzed using custom software (Labview and Igor Pro, respectively) and used to determine the rupture force, which was marked as the maximum force sustained by the attachment during each event. All the individual rupture force values and calculated mean rupture strengths are provided in Table S2.

---

**Supplementary Table 1.**
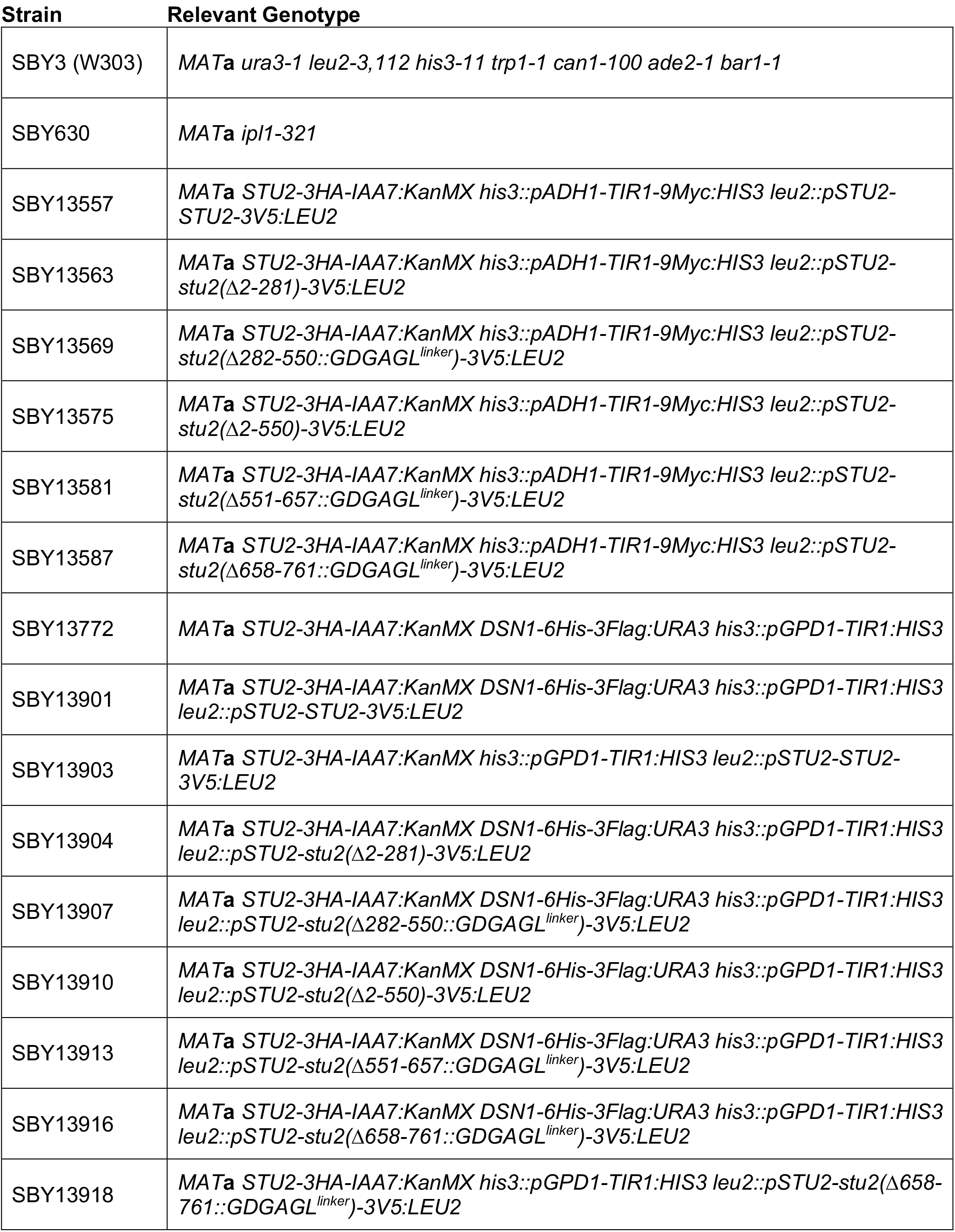
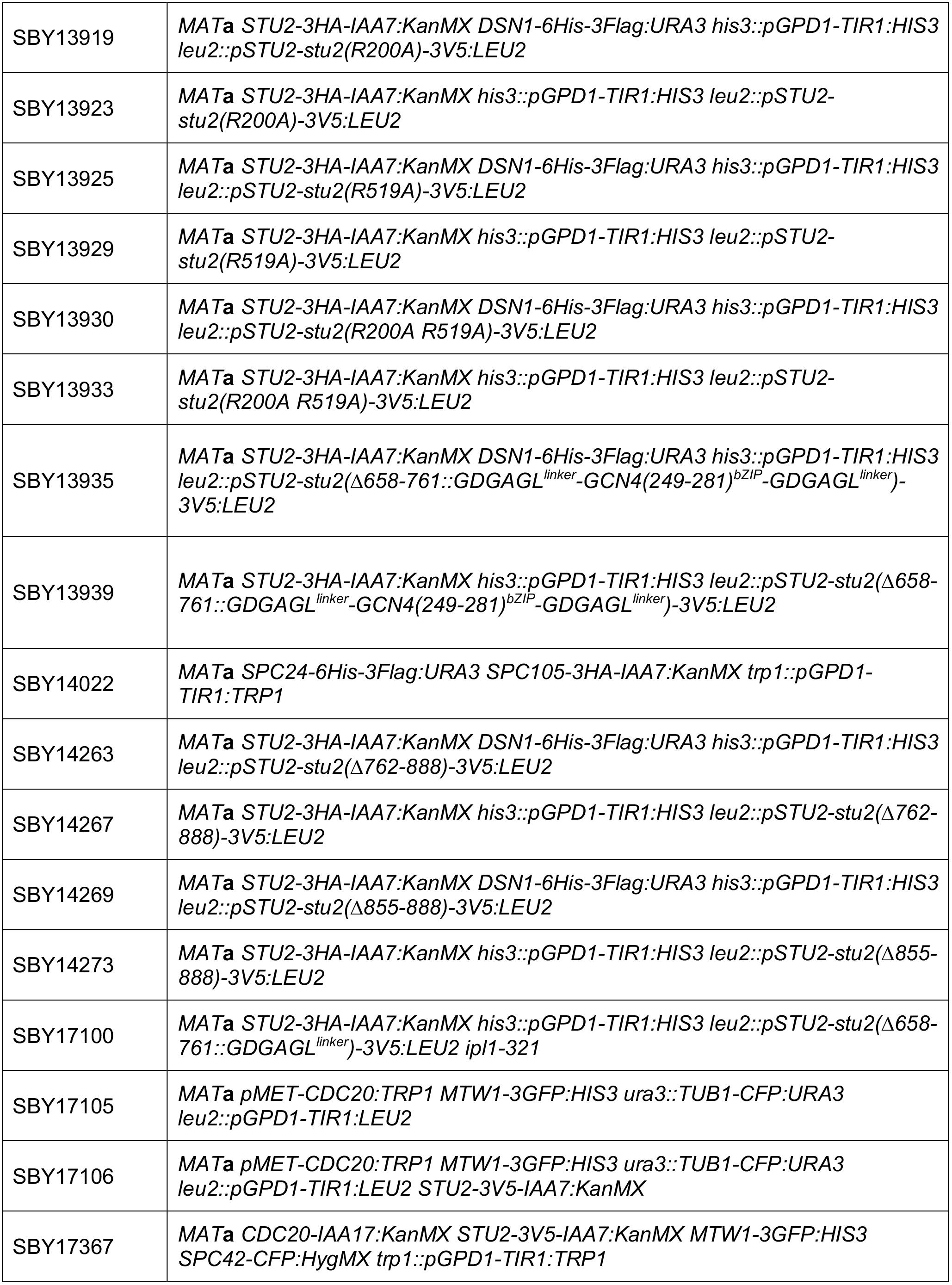
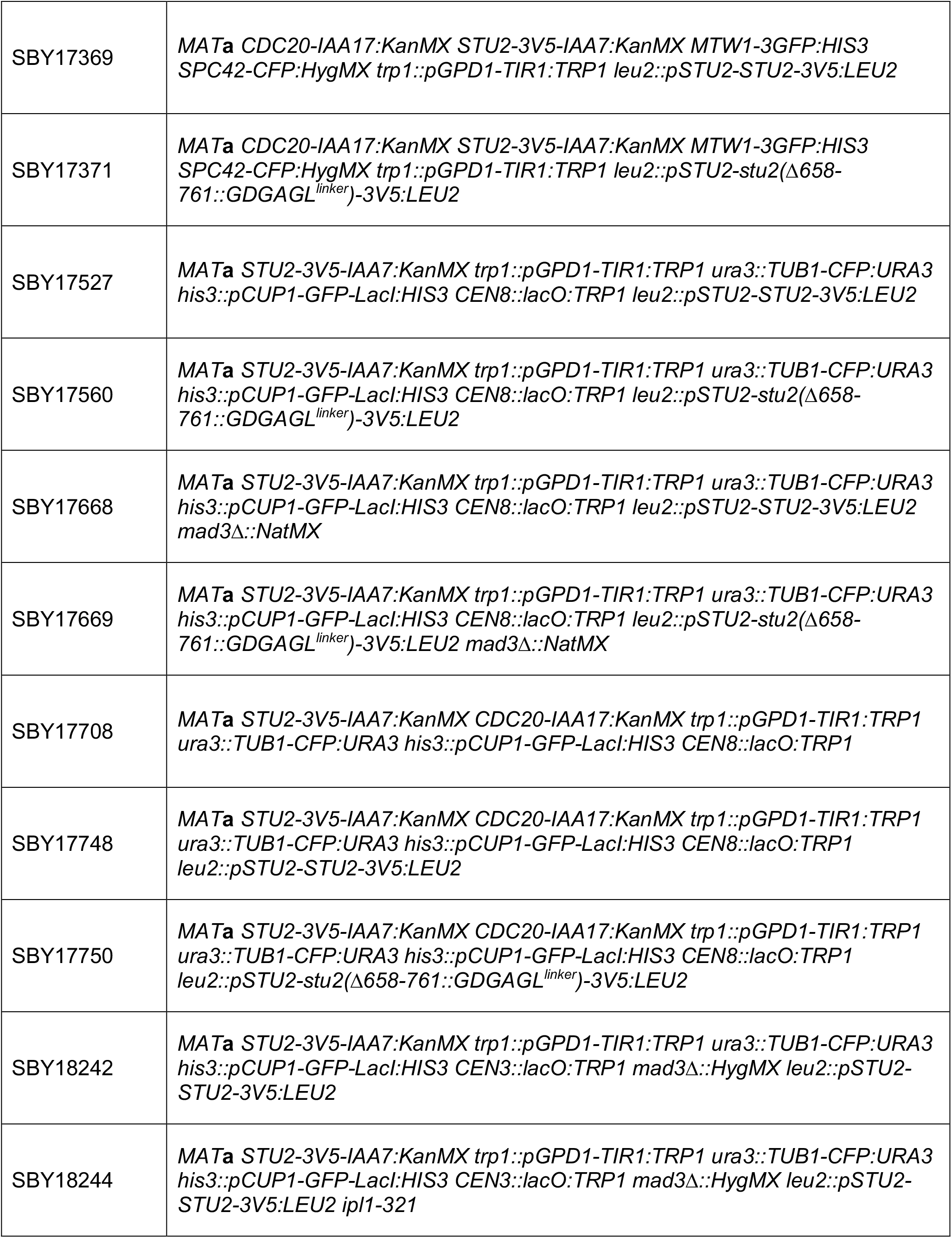
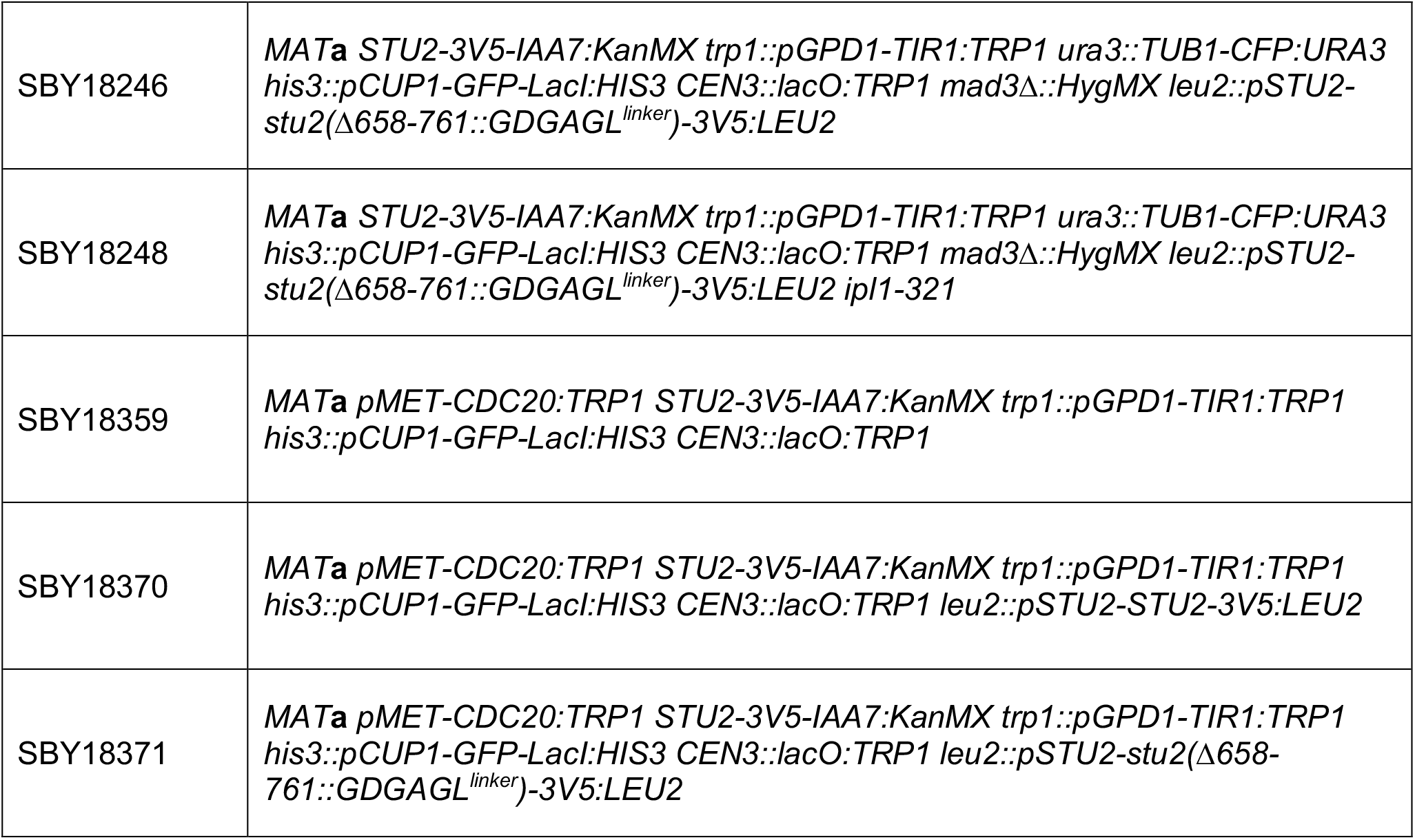
Strains used in this study. All strains are derivatives of SBY3 (W303)

**Table S2.**
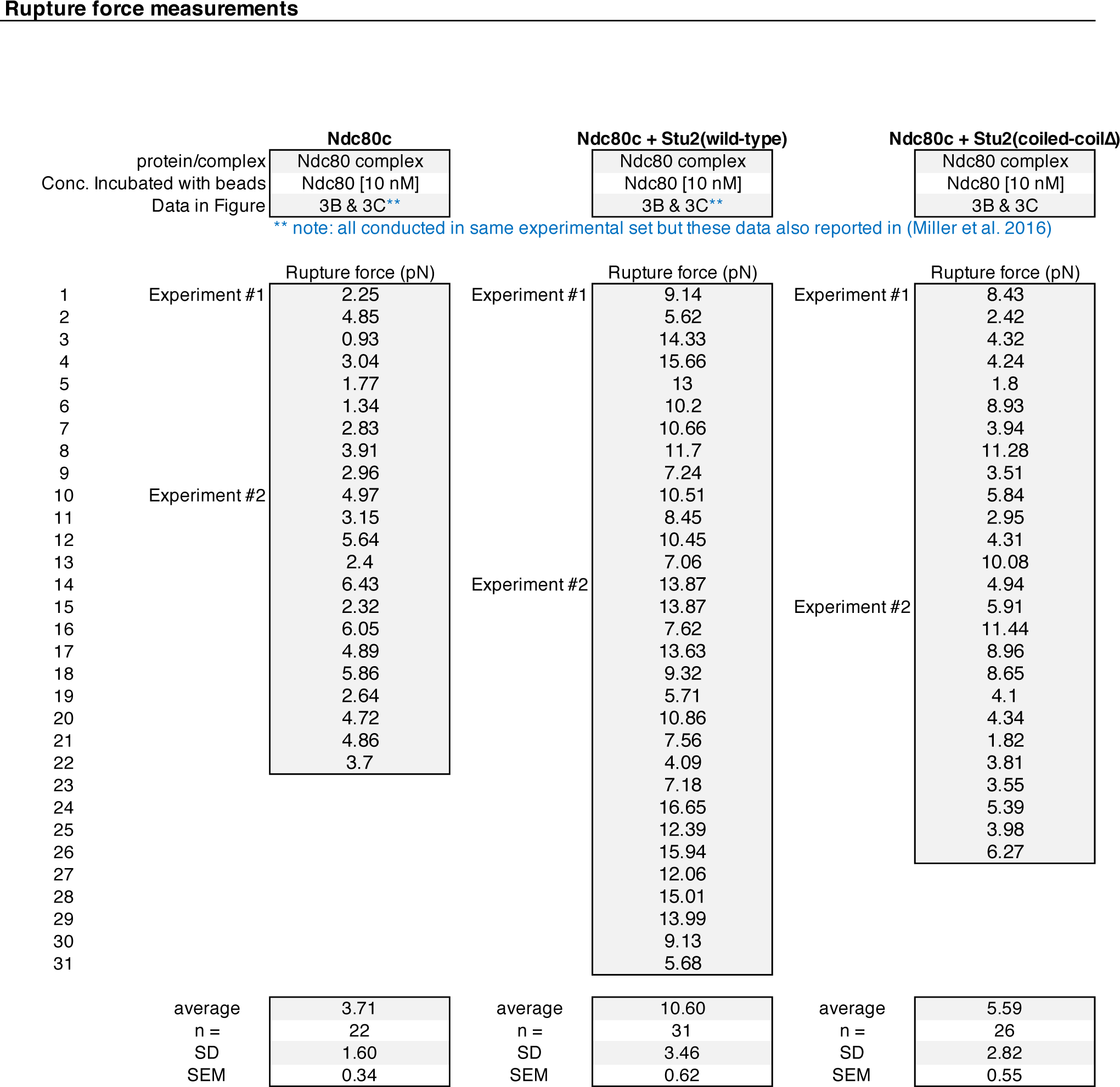
Summary of optical trap-based bead motility assays, related to Figure 2

